# Heritable changes in division speed accompany the diversification of single T cell fate

**DOI:** 10.1101/2021.07.28.454102

**Authors:** Marten Plambeck, Atefeh Kazeroonian, Dirk Loeffler, Timm Schroeder, Dirk H. Busch, Michael Flossdorf, Veit R. Buchholz

## Abstract

Rapid clonal expansion of antigen specific T cells is a fundamental feature of adaptive immune responses. It enables the outgrowth of an individual T cell into thousands of clonal descendants that diversify into short-lived effectors and long-lived memory cells. Clonal expansion is thought to be programmed upon priming of a single naïve T cell and then executed by homogenously fast divisions of all of its descendants. However, the actual speed of cell divisions in such an emerging ‘T cell family’ has never been measured with single-cell resolution. Here, we utilize continuous live-cell imaging *in vitro* to track the division speed and genealogical connections of all descendants derived from a single naïve CD8^+^ T cell throughout up to ten divisions of activation-induced proliferation. This comprehensive mapping of T cell family trees identifies a short burst phase, in which division speed is homogenously fast and maintained independent of external cytokine availability or continued T cell receptor stimulation. Thereafter, however, division speed diversifies and model-based computational analysis using a novel Bayesian inference framework for tree-structured data reveals a segregation into heritably fast and slow dividing branches. This diversification of division speed is preceded already during the burst phase by variable expression of the interleukin-2 receptor alpha chain. Later it is accompanied by selective expression of memory marker CD62L in slower dividing branches. Taken together, these data demonstrate that T cell clonal expansion is structured into subsequent burst and diversification phases the latter of which coincides with specification of memory vs. effector fate.

**Significance:** Rapid clonal expansion of antigen-specific T cells is a fundamental feature of adaptive immune responses. Here, we utilize continuous live-cell imaging *in vitro* to track the division speed and genealogical connections of all descendants derived from a single naïve CD8^+^ T cell throughout up to ten divisions of activation-induced proliferation. Bayesian inference of tree-structured data reveals that clonal expansion is divided into a homogenously fast burst phase encompassing two to three divisions and a subsequent diversification phase during which T cells segregate into quickly dividing effector T cells and more slowly cycling memory precursors. Our work highlights cell cycle speed as a major heritable property that is regulated in parallel to key lineage decisions of activated T cells.

## Introduction

The smallest unit from which an adaptive immune response can originate is an individual antigenspecific lymphocyte (1). For both CD4^+^ and CD8^+^ T cells, it has been shown that single-cell-derived immune responses *in vivo* are subject to immense variation, despite being directed against the same epitope and unfolding within the same host (2–6). Upon vaccination or infection even naïve T cells harboring identical T cell receptors (TCRs) will generate ‘T cell families’ (i.e. immune responses derived from a single T cell) of highly distinct size and phenotypic composition (2, 3, 6). Interestingly, within a given T cell family, clonal expansion and T cell differentiation are interdependent: At the peak of expansion, larger T cell families harbor lower percentages of long-lived central memory precursors (CMPs) and higher percentages of shorter-lived effector memory precursors (EMPs) and terminal effectors (TEs) than smaller T cell families (2, 7).

To account for the variation in single-cell-derived expansion and the interdependency of T cell phenotype and family size, we have put forward a stochastic developmental framework in which naïve T cells first give rise to slowly dividing CMPs, which can then differentiate into more quickly dividing but shorter-lived progeny (2). This framework proved to be efficient in describing features of singlecell-derived T cell responses *in vivo*, such as the relative independence of a T cell family’s memory capacity from its acute size of clonal expansion. This framework is further supported by direct measurements of division speed *in vivo*, showing that, already by day four after vaccination, CD62L^+^ CMPs undergo on average one division less per day than their CD62L^-^ counterparts (8). Moreover, recent work confirmed the largely unidirectional differentiation of CMPs into non-CMPs during the expansion phase of a T cell response (9) and the role of CMPs as the major source of long-lasting CD8^+^ T cell memory (10–12).

However, certain features of our originally proposed stochastic framework are at odds with observations made during the very early phase of T cell activation: First, directly after activation *in vivo*, CD8^+^ T cells have been found to divide particularly fast (13, 14), conflicting with the idea of an initial emergence of slowly-dividing CMPs. Second, elegant *in vitro* experiments have shown that division activity is strongly correlated among the members of a given T or B cell family, arguing against the emergence of distinct expansion kinetics within the same family (15–18). However, these latter studies mainly investigated the first three to four cell divisions executed by an expanding lymphocyte family and compared division speed only between close relatives within the family tree (i.e., sibling or mother-daughter correlations). We reasoned that to investigate the gradual cross-generational emergence of slower- or faster-dividing genealogical branches, division speed and T cell kinship must be tracked across longer genealogical distances than done before.

Therefore, we performed continuous *in vitro* imaging of single-cell-derived clonal expansion for up to five days after T cell activation. In contrast to previous studies, we utilized a culture system that allowed us to faithfully track the genealogical connections within expanding T cell families for up to ten generations. We found that after completing their first cell cycle, CD8^+^ T cells underwent a burst phase of two to three uniformly quick divisions. Mean division speed then slowed down in absence of continued TCR stimulation, and remained high when TCR stimulation was maintained. However, even upon continuous TCR stimulation, distinctly proliferating branches emerged in the later generations of a family tree. To better quantify the hereditary nature of this process, we developed a computational framework that enabled a model-based analysis of the tree-structured data obtained from live-cell imaging. We combined branching process modeling with a Bayesian inference approach for models with hidden states. This framework enabled us to test various hypotheses about the diversification pattern of proliferating T cells into subsets with distinct inter-division times. These analyses identified a model in which naïve T cells first differentiate into a quickly proliferating early-activated state, which then transits into slowly and quickly dividing branches that heritably maintain their distinct division activity. Further investigating the molecular regulation of these processes, we found that distinct expression levels of the high affinity interleukin-2 receptor alpha chain (CD25), established during the burst phase, preceded the adoption of distinct division activities. Moreover, we found that CD62L, as a marker of CMP differentiation, was preferentially expressed in slowly dividing T cells that emerged beyond the burst phase.

Taken together, our work shows that after a short burst phase the division speed of activated CD8^+^ T cells segregates into slow and fast cycling branches. Moreover, it provides novel mathematical methodology suited to test models of heredity within tree-structure data, generated by an expanding T cell family or any other proliferating and continuously-imaged cell type.

## Results

### Continuous *in vitro* imaging reveals that distinct division speeds emerge within the same T cell family

In order to comprehensively map clonal expansion *in vitro* starting out from individual naïve CD8^+^ T cells, we utilized a continuous imaging platform (19, 20). Naïve CD44^low^ CD8^+^ T cells were individually sorted via flow cytometry from peripheral blood of C57BL/6 mice, transferred to culture wells coated with anti-CD3 and anti-CD28 antibodies and supplemented with interleukin 2 (IL-2) **(Fig. 1 A)**. This treatment enabled TCR restimulation throughout the expansion phase and induced vigorous proliferation that we monitored via continuous imaging for up to five days **(Fig. 1 B and Movie S1)**. Importantly, IL-2 concentrations remained at saturating levels throughout the whole observation period **(Fig. S1)**. Since overcrowding of microwells (20-100 μm in diameter) and clustering of T cells can be a problem for tracking the individual members of an expanding T cell family tree (16, 17), we sorted single T cells into relatively large wells with a diameter of 1840 μm and imaged the complete well **(Movie S2)**. These “macrowells” allowed for freer migration and reduced overlay phenomena of activated T cells. This enabled reliable tracking of individual T cells for up to 10 cell generations **(Fig. 1C-D and Movie S3)**. We found that activated T cells required on average 39 hours to complete their first cell division **(Fig. 1 E)**. The average interdivision time for subsequent cell cycles amounted to 8.6 hours but showed strong variation ranging from 5 to 28 hours per cell cycle **(Fig. 1 F)**. Interestingly, while some of this variation could be attributed to differences between the overall division speed of distinct T cell families (interfamily variance), most of it was derived from differences in division speed between individual T cells belonging to the same family (intrafamily variance) **(Fig. 1G)**. To explore whether these differences arose as random fluctuations or were heritably maintained, we first investigated correlations of division speed between close relatives in a family tree. As previously reported (14–17), we found that T cell siblings shared similar division speeds with one another and with the mother cell from which they were derived **(Fig.1 D and H)**. To then investigate more distant genealogical relationships, we grouped the progeny of an individual T cell into four main branches, emerging from the second generation of each family tree **(Fig. 1 I)**. In more than 40% of family trees (18 of 43), the average T cell division speed differed significantly between these branches **(Fig. 1 J)**. We also identified distinct average division speeds in branches derived from the first generation of T cell family trees, albeit at a lower incidence **(Fig. S2)**. These data show that slow vs. fast cycling T cells can segregate onto distinct branches of the same T cell family tree. To more formally explore whether this segregation could be accounted for by differentiation of proliferating T cells into heritably distinct subsets we next performed computational modelling.

**Figure 1:**
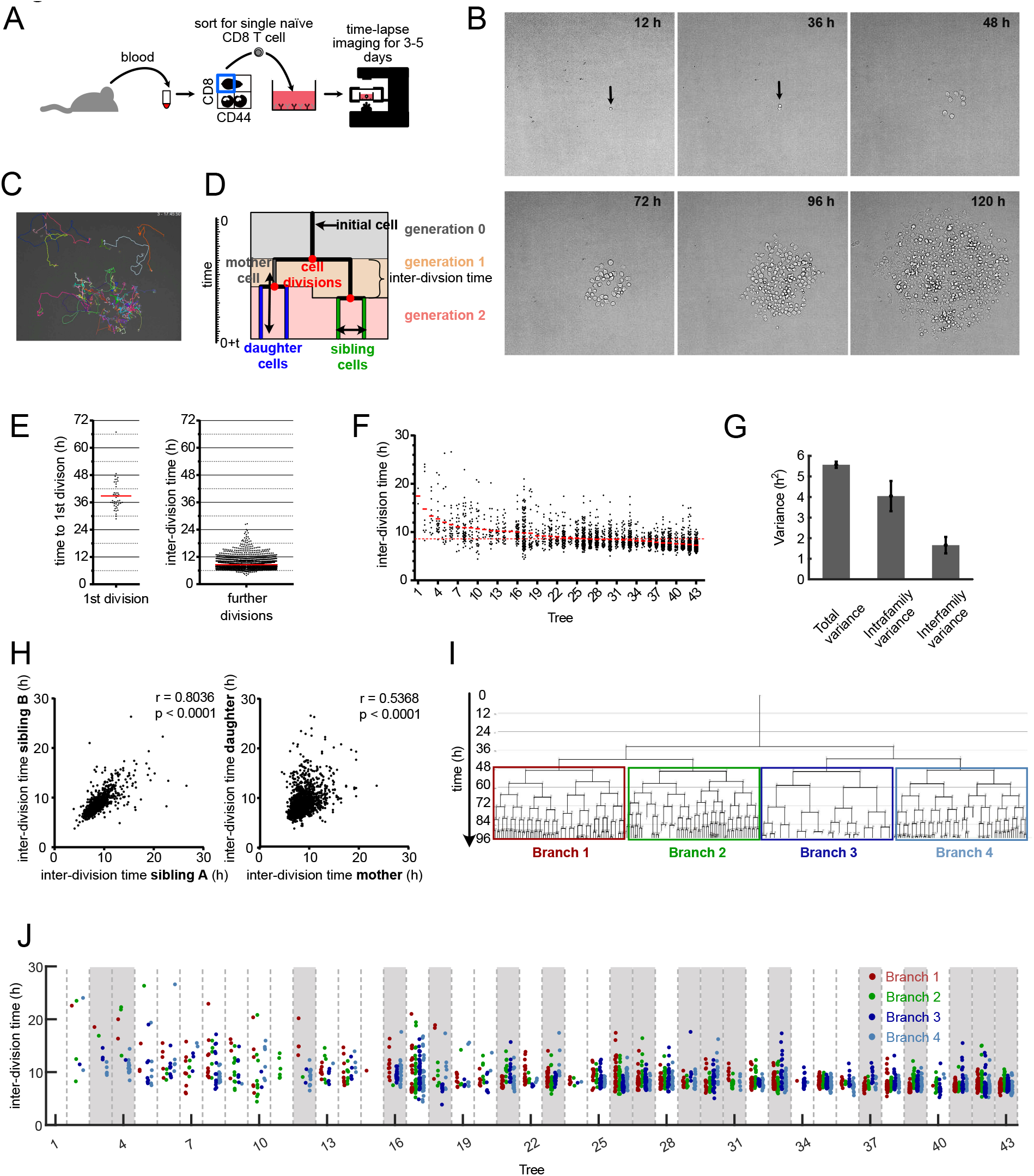
Continuous *in vitro* imaging reveals that distinct cell cycle speeds emerge within the same T cell family. **(A)** Blood was taken from an OT-I mouse and sorted for naïve CD8+, CD44^low^ cells. A single naïve OT-I cell was sorted into a well that was coated with αCD3 and αCD28 in the presence of 25 U/mL IL-2. The cell culture plate was transferred to a live-cell imaging microscope and imaged for the next three to five days. **(B)** Pictures, taken at different time points from the same single cell-derived progeny. **(C)** Snapshot of a single cell-derived progeny at 3 days, 17 hours and 45 minutes after start of acquisition. Colored lines represent migration pattern of individual cells. **(D)** Definitions of family tree associated data: y-axis: Time. Generation: The number of cell divisions that have occurred from the naïve T cell until the respective cell was created by division of its respective mother cell. Mother and daughter cells: The two cells that originate from the same cell division are daughter cells in respect of the original cell that divided (mother cell). Sibling cells: Cells that have the same mother cell. Interdivision time: Time between the creation of an individual cell due to the division of its mother cell into two daughter cells and the end of the respective cell due to its own division into two new daughter cells. **(E)** Time for first and subsequent divisions. Single naïve (CD44^low^) CD8^+^ T cell were sorted into separate wells of an αCD3/CD28-coated 384-well plate and imaged for 5 days in a live-cell imaging microscope. The fate of each cell was tracked and the inter-division time for the first division and all subsequent divisions were determined for 43 single-cell derived progenies (in total 43 cells for 1^st^ division and 2710 cells for subsequent cell divisions). Red lines indicate the means. **(F)** Intra- and inter-clonal variability of inter-division times. The inter-division times from (E) (first division excluded) arranged according to their mean inter-division time. Red bars: mean inter-division time within family tree. Red dotted line: mean for all cells. **(G)** Total variance of inter-division times from further divisions in (E), as well as the contribution of intrafamily and interfamily variances are shown. Intrafamily variance is calculated as the weighted mean of the variances of inter-division times within all different families. Interfamily variance is calculated as the weighted variance of the mean inter-division times of different families (Supplementary Methods). Total variance: 5.57±0.15; intrafamily variance: 4.045±0.73; interfamily variance: 1.66±0.39. The distribution of intrafamily variance is significantly larger than that of interfamily variance (p-value < 1e-3). **(H)** Inter-division times from (E) were plotted against the interdivision times of their sibling cell (left panel, 1201 pairs) or their direct daughter cells (right panel, 2630 pairs). The correlation coefficient r and the p-value for spearman correlation are indicated. **(I)** A representative family tree was divided into 4 branches starting from generation 2. Loose ends: no further tracking possible for the respective cell. **(J)** Inter-division time of cells in the four branches starting from the second generation—as depicted in (I)—are color-coded for all trees in (F). In 18 out of 43 trees (~ 42%, trees highlighted in gray), inter-division times differed significantly between the four branches. For every tree, a p-value was calculated based on one-way ANOVA. We then estimated positive false discovery rates (pFDR) for multiple hypothesis testing based on these p-values, and used a cutoff of pFDR<0.05 for significance.

### Computational analysis of T cell family trees identifies a model in which early activated T cells diverge into slow- and fast-cycling subsets

To test whether differentiation of proliferating T cells into subsets with heritably distinct division speeds indeed accounted for the variability of inter-division times and the resulting interior structure of T cell family trees, we used a branching process framework (21). Since we couldn’t monitor the underlying differentiation processes directly, we modeled the differentiation state of each cell as a hidden variable (i.e. its belonging to a slow- or fast-cycling subset). To infer the parameters of the branching process, we then developed a Bayesian inference framework for tree-structured data, comprised of a Markov Chain Monte Carlo (MCMC) sampling approach with hidden layers (22, 23). Our inference framework takes as input the genealogical trees obtained from live cell imaging movies, as well as a model hypothesis describing the underlying branching process. As output, it returns the posterior distribution of model parameters as well as the model evidence for every assumed model hypothesis. These model evidences and corresponding Bayes factors are then used for model selection (**Fig. 2 A and Supplementary Methods**). The individual steps of our iterative scheme are depicted in **Figure 2 B** (**Fig. 2 B and Supplementary Methods**).

**Figure 2:**
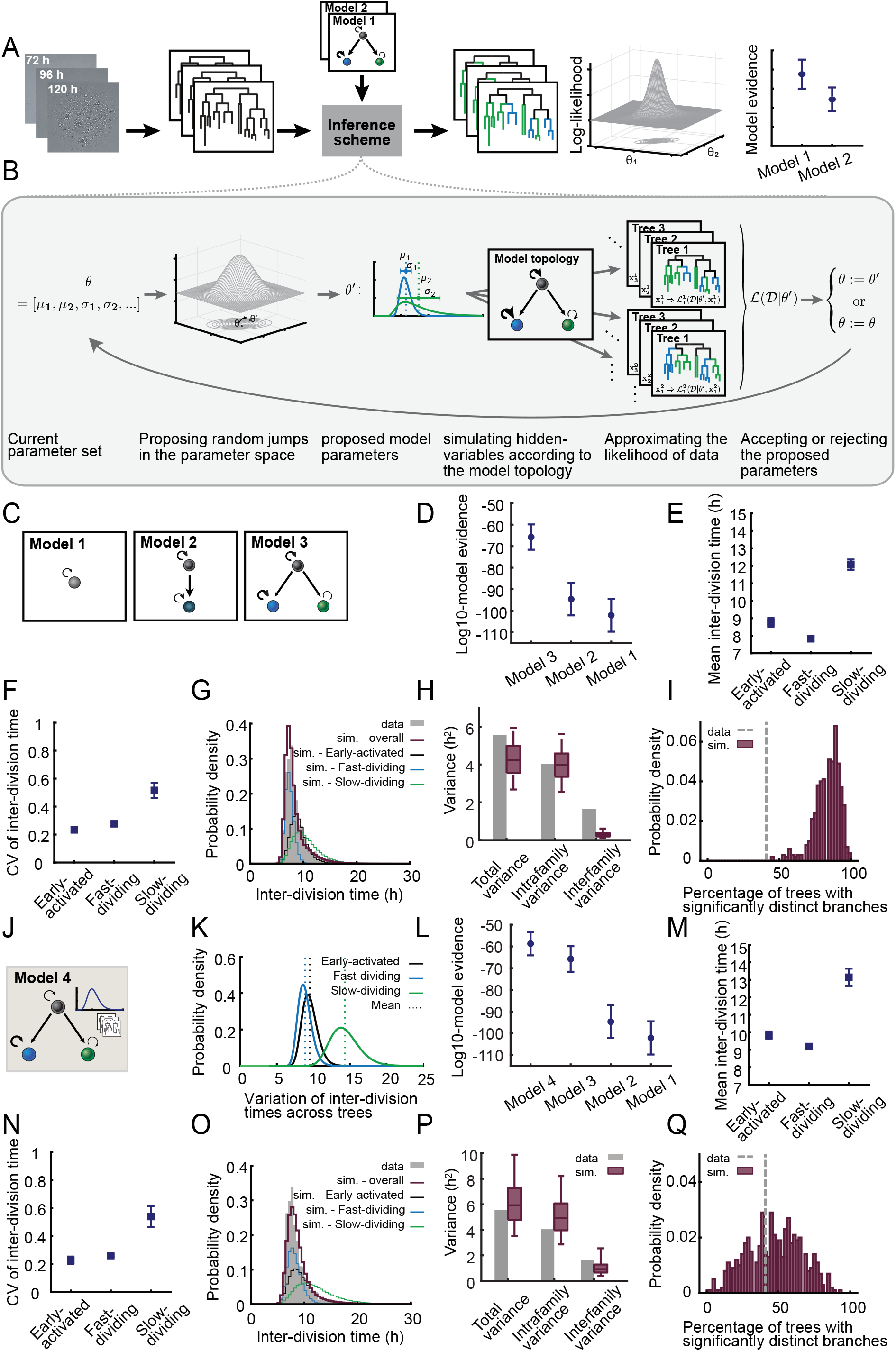
Computational analysis of T cell family trees identifies a model in which early activated T cells diverge into slow and fast cycling subsets. **(A)** Schematic of the workflow of our inference scheme. The scheme requires as input 1) lineage trees obtained from successive 2D microscopy images, and 2) one or more model hypotheses describing the underlying branching process. For every model hypothesis, it returns posterior distribution of model parameters and an approximation of the model evidence. The model evidences are used for model selection. **(B)** Individual steps of the iterative inference scheme. In every iteration, a new set of model parameters are proposed based on the current parameter values and a proposal distribution. These proposed parameters, e.g., the mean and CV of subset-specific inter-division time distribution, are used to simulate several samples of hidden variables of the model. For instance, the shown model topology includes three different subsets (black, blue and green). The simulated hidden variables in this case are the subset that is assigned to each cell in the tree. For every tree and every simulated sample, the likelihood of the data is calculated given the current value of parameters and hidden variables. A Monte Caro approximation method is used to calculate the overall likelihood of the data given the parameters. The proposed parameter set is accepted with the acceptance probability calculated based on the likelihood, and is rejected otherwise. **(C)** Schematic of different model hypotheses used for the analysis of T cell family trees: Model #1 assumes a homogenous population with one proliferation speed. Model #2 assumes an early expanding subset that later differentiates into another subset with a distinct proliferation speed. Model #3 assumes an early expanding subset that diversifies into slow- and fast-dividing subsets. **(D)** The model evidences for the models in (C) indicate the following hierarchy: model #3 > model #2 > model #1; model #3, having the highest evidence, explains the data best and model #1, having the lowest evidence, is least matching to the data. The circles show the mean of the base-10 logarithm of model evidences of the eight data groups (Supplementary Methods). The error bars show the standard error of these means. **(E)** Inferred mean and **(F)** CV of the inter-division time distribution for “Early-activated”, “Slow-dividing” and “Fast-dividing” subsets based on model #3. For every data group, the median of the posterior distribution of the respective parameters is calculated; the squares denote the mean of these medians (Supplementary Methods). The error bars show the standard error of the mean. **(G)** The distribution of inter-division times in 10000 simulated trees based on model #3 compared to that of the experimental data (grey). The red histogram shows the overall distribution in the simulated data, while the black, blue and green histograms show the distribution of “Early-activated”, “Fast-dividing” and “Slow-dividing” subsets respectively. The results are shown for one data group. **(H)** Total variance of the inter-division times and the contribution of intrafamily and interfamily sources as observed in the experimental data (grey bars) and simulated data (red boxes). Intrafamily variance is calculated as the weighted mean of the variances of inter-division times within different families. Interfamily variance is calculated as the weighted variance of the family mean inter-division times (Supplementary Methods). The simulated data consists of 500 datasets of each 44 trees simulated based on model #3. The boxes show the 90%-confidence interval of the simulated data. The total variance and intrafamily variance of the experimental data are contained within the 90%-confidence interval of the simulated data, while for interfamily variance, the simulated data does not contain the experimental data point. The three variance terms from left to right for the experimental data: 5.57, 4.05, 1.66. The 5-percentile, median and 95-percentile for the three variance terms for simulated data from left to right: [2.68, 4.23, 5.93], [2.56, 3.98, 5.61], [0.11, 0.27, 0.61]. The results are shown for one data group. **(I)** Percentage of the trees whose four branches (as in Fig. 1I) have significantly distinct inter-division times. The grey line shows the experimental data and the red histogram shows the distribution of this percentage in the simulated data as in (H) (Supplementary Methods). The results are shown for one data group. **(J)** Model #4 adopts the topology of model #3 and incorporates variability between trees in addition. The mean inter-division times of subsets EA, S, and F are assumed to differ between trees and this variation is assumed to be log-normally distributed. The inset shows the stack of different trees and a sample log-normal distribution for the variation of subset-specific mean inter-division times between these trees. (Supplementary Methods). **(K)** Inferred variation of subset-specific mean inter-division times among different families based on model #4. The black, blue and green curves show the distribution for “Early-activated”, “Fast-dividing” and “Slow-dividing” subsets respectively. The results are shown for one data group. **(L)** Model evidences for the models in (C) and (J) indicate that model #4 explains the data best. The circles show the mean of the base-10 logarithm of model evidences of eight data groups (Supplementary Methods). The error bars show the standard error of the mean. A difference of seven between the mean log10-model evidences for model #4 and model #3 indicates the significant superiority of model #4. The mean Bayes factor, as well as individual Bayes factors in every data group, show at least a strong evidence for choosing model #4 over model #3 (Fig. S5 and S12). **(M)** and **(N)** Same as (E) and (F) with parameters inferred based on model #4. **(O), (P)** and **(Q)** Same as (G), (H) and (I) with simulated data based on model #4. **(P)** The total variance, intrafamily variance and interfamily variance of the experimental data are all contained within the 90%-confidence interval of the simulated data. The three variance terms from left to right for the experimental data: 5.57, 4.045, 1.66. The 5-percentile, median and 95-percentile for the three variance terms for simulated data from left to right: [3.5, 5.92, 9.89], [2.86, 4.92, 8.21], [0.39, 0.91, 2.55].

We assumed that the inter-division times of cells belonging to every subset are distributed according to a log-normal distribution with a specific mean and coefficient of variation (CV). To allow for efficient computation we divided our dataset into eight groups of five family trees each, and for each group, inferred the posterior distribution of model parameters as well as the model evidences. We then selected the models fitting best to our data by calculating Bayes factors based on the model evidences. To further assess how well the best-fitting model represented our data, we tested whether model-based simulations recapitulated the statistical characteristics of the experimental data (**Supplementary Methods**).

We first investigated three basic model topologies: In model #1 naïve T cells gave rise to one proliferating subset, in model #2 naïve T cells gave rise to one proliferating subset that could differentiate into another subset proliferating at a distinct speed, and in model #3 naïve T cells gave rise to one proliferating subset that could differentiate into two others each proliferating at distinct speeds **(Fig. 2 C)**. As expected, model #1, which assumed that proliferation of all activated T cells could be described by one division speed (i.e., one distribution of division speed learned from the data) was insufficient account for the specific structure of T cell family trees (model evidences and Bayes factors shown in **Fig. 2 D and S3-6)**. Interestingly, model #2, which relied on the differentiation of one proliferating subset into another, was also insufficient to account for the measured data. Only model #3, which assumed that a proliferating early-activated subset differentiates into a slowly or a quickly dividing one, provided an adequate fit to the measured data **(Fig. 2 D and S3-6)**. Mean inter-division times of these subsets were predicted at 8.8 h, 7.8 h and 12 h for the early-activated (EA), fast-dividing (F) and slow-dividing (S) subsets, respectively **(Fig. 2 E and S7)**. The learned distribution of division speed for the slow-dividing subset was relatively wide, while those of the early-activated and fast-dividing subsets showed considerably less variation **(Fig. 2 F and S7)**. To rule out the possibility that model #3 was the preferred model only due to its additional flexibility, we compared it to a mixture model in which, once leaving the EA state, cell cycle speeds are taken from two overlaid distributions without considering the topological restraints of the proposed differentiation process **(Supplementary Methods)**. The mixture model could not explain the experimental data as well as model #3, since it did not consider the topological relationship in T cell family trees, which are accounted for in model #3 (the model evidences and Bayes factors are shown in **Fig. S3-6)**. Thus, the topological diversification of cells into distinct subsets and the maintenance of distinct division speeds within every subset, as considered in model #3, were crucial for explaining the data. Using the simulation analysis mentioned earlier, we found that the best-fit “EA to S or F” model (model #3)—although adequately reproducing the overall distribution of inter-division times **(Fig. 2 G and S8)**—underestimated the contribution of interfamily differences to the overall variation of T cell division speed **(Fig. 2 H and S9)**. It further overestimated the percentage of trees with branches showing significantly distinct division speeds **(Fig. 2 I and S10)**. This indicated that division speed of activated T cells may not only depend on their current differentiation state (EA, S or F) but also on an ancestral imprinting received by the starting cell and passed down to its descendants. To account for this clonal imprinting of division speed, we allowed for a log-normally distributed factor in the model formalism that could modify all subset-specific proliferation rates (for subsets EA, S and F) of a given T cell family by a certain value **(Supplementary Methods)**. Fitting this extended model (model #4, **Fig. 2 J)** to the data, we quantified the variation of subset-specific mean inter-division times between different families **(Fig. 2 K and S11)** and found that this model was best supported by the data. A difference of seven in the average log10-model evidences indicated that model #4 was significantly better than model #3 in explaining the data **(Fig. 2 L)**. Furthermore, the corresponding Bayes factors showed that in six out of eight data groups, there was a decisive evidence for choosing model #4 over model #3 (log10-Bayes factors > 2), with the remaining two data groups providing a strong evidence for model #4 (log10-Bayes factors > 1) **(Fig. S3-5 and S12)**. Importantly, despite the additional variation introduced by this factor, the distinct division speeds of the EA, F and S subsets were maintained according to the best-fit parameters of model #4 **(Fig. 2 M and N and S13)**. We noted that the overall distribution of inter-division times in simulated data based on model #4 **(Fig. 2 O and S14)** was similar to the results of model #3 **(Fig. 2 G and S8)**, and this statistical feature alone would not have been enough to distinguish between the two models. Instead, more elaborate structural features of the T cell family trees could demonstrate differences between the two models. Indeed, model #4 now correctly captured both intra- and inter-family variability **(Fig. 2 P, S15)** and generated a realistic fraction of T cell family trees whose second-generation branches proliferated at distinct average speeds **(Fig. 2 Q, S16)**. The inferred model parameters and model evidences for all models and data-groups are depicted in **Supplementary Figures S7, S13 and S17-S19**. Taken together, these computational analyses suggested that T cell clonal expansion is not a homogenous process programmed exclusively upon T cell priming. Instead, the emergence of multiple T cell subsets proliferating at distinct speeds appeared necessary to correctly capture the evolution of an expanding T cell family tree.

### Diversification of division speed occurs after a homogenous burst phase and is preceded by differences in CD25 expression

The experimental conditions, under which the above-mentioned results were gathered, allowed both the starting cell as well as its descendants to receive TCR stimulation. This raised the question whether the emergence of slow and fast dividing branches within the same T cell family tree was the consequence of TCR stimuli incidentally accumulating in one branch but not the other. Thus, we next asked whether distinct division speeds would also emerge when TCR stimuli are restricted to the starting cell. To achieve this, we activated T cells in bulk via plate bound anti-CD3 and anti-CD28 antibodies in presence of IL-2 and IL-12 and, 24 h later, sorted single undivided T cells into wells containing no further TCR stimuli. To sustain proliferation for an extended period of time, wells were supplemented with saturating doses of IL-2 only (“brief”) or IL-2 and IL-12 (“brief + IL-12”). Since IL-12 induces the expression of the high affinity IL-2 receptor alpha chain (CD25) we were expecting a synergistic effect of both cytokines jointly added. As before we tracked T cell proliferation via continuous imaging for up to five days **(Fig. 3 A)**. Upon brief TCR stimulation, 1^st^ and 2^nd^ generation T cells divided at the same average speed as those cultured in the sustained presence of TCR stimuli (“sustained”). After this initial burst phase, the average division speed of briefly stimulated T cells significantly slowed down, albeit less in the group supplemented with IL-2 and IL-12 **(Fig. 3 B and C)**. Importantly, the variation in division speed of both the “brief” and “brief+IL-12” groups matched or even exceeded that of T cells receiving sustained TCR stimuli **(Fig. 3 B and C)**. If this variation arose due to random fluctuations in division speed, one would expect that, over time short and long inter-division times cancel each other out and all members of a growing T cell family occupy the same or adjacent generations. However, we found that upon acquiring an increasing number of divisions, the generational range between the most and least divided members of a T cell family constantly widened (increasing length of colored rows from 36 to 96h), arguing in favor of distinct family branches heritably maintaining slower and faster cell cycle activity **(Fig. 3 D)**. To more closely investigate the mechanistic origin of these distinct division speeds, we measured the expression of the high-affinity IL-2 receptor alpha chain (CD25) during continuous imaging. We achieved this by addition of very low concentrations of anti-CD25 antibody conjugated to relatively photostable fluorophores such as Phycoerythrin (PE) or Allophycocyanin (APC) (20, 24). We found that addition of very low antibody concentrations had negligible effects on proliferative activity of activated T cells **(Fig. S20)** while generating robust and reliable fluorescent signals **(Movie S4)**. Interestingly, T cells receiving brief vs. sustained TCR stimulation showed distinct levels of CD25 surface expression already within the 1^st^ and 2^nd^ generation but still proliferated at the same speed. Only from the 4^th^ generation onwards did lower or higher CD25 expression levels begin to correlate with lower or higher T cell division speed **(Fig. 3 E)**. In fact, closer inspection of individual T cell family trees showed that changes in CD25 expression levels can be allocated to specific branches and can precede changes in division speed that develop across multiple generations **(Fig. 3 F and Fig. S21)**. Taken together, these data indicate that during the first two generations of T cell proliferation, division speed is largely independent of sustained TCR stimuli, IL-12 availability or CD25 expression levels. Thereafter, however, distinct levels of CD25 surface expression begin to correlate with distinct division speeds and are heritably maintained within distinct branches of an expanding family tree.

**Figure 3:**
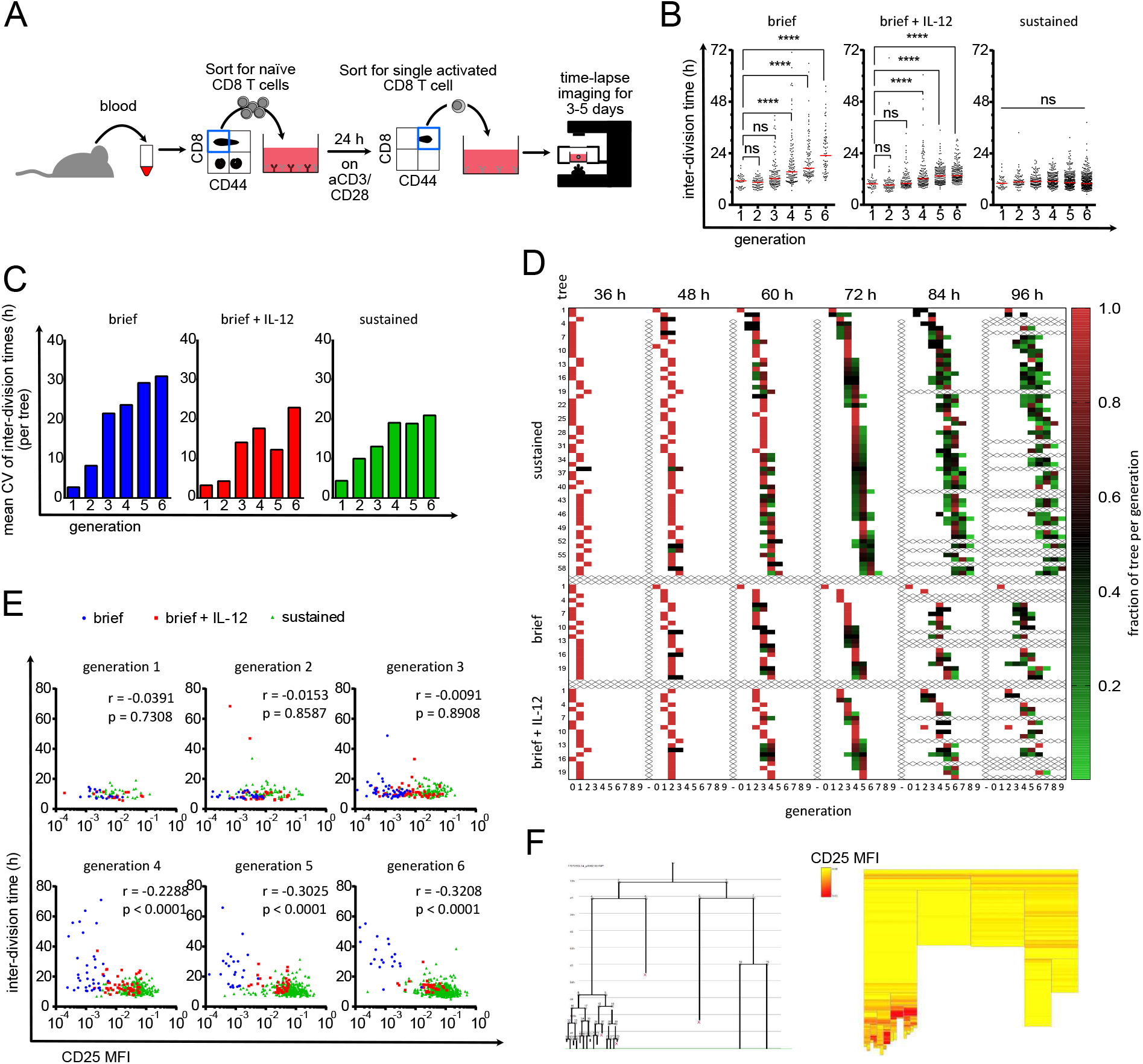
Diversification of division speed occurs after a homogenous burst phase and is preceded by differences in CD25 expression. **(A)** Live-cell imaging with brief TCR stimulation. Blood was taken from an OT-I mouse and sorted for naïve CD8 T cells. 10 000 cells/well were activated for 24 h with plate-bound αCD3/CD28 and 25 U/mL IL-2. Cells were sorted again for activated (CD44high) cells and a single cell was sorted in each well of a 384-well plate that was coated with ICAM-1 or αCD28 to enable attachment. The cells were imaged for 3-5 days. **(B)** Cells were stimulated as described in (A). After the brief stimulation and segregation 10 ng/mL IL-12 were added to the medium (second panel) or not (left panel). The cells in the right-hand panel were stimulated continuously with αCD3/αCD28 as in figure 1. The inter-division times of all cells are plotted for the respective generations. Red lines indicate the median. Kruskall-Wallis test: ****: p<0.0001 **(C)** Data points from (B) were allocated to the different family trees, and the coefficients of variation of the inter-division times within the trees were calculated within each generation. The mean of these CVs is plotted as bar graph. On average 15.45 trees per bar (4–27). **(D)** For all three conditions it is shown how the individual cells within a tree are distributed over the generations at different time points. The fraction of tree per generation is calculated by dividing the number of cells in a specific generation at the given time point by the maximum number of cells potentially present in the respective generation (number of cells in generation X / 2 ^generation^ X). E.g., for a potential tree that consists after 48 h out of 1 cell in generation 2 and 6 cells in generation 3 the respective fractions of tree per generation are for generation 2: 1 / 2^2^ = 0.25 and for generation 3: 6 / 2^3^ = 0.75. For incompletely tracked trees (e.g., when cells died or the identity of cells is unclear) the fractions of tree per generation is corrected so that the sum of all fractions of tree per generation sum up to 1.0 (corrected fraction of tree per generation = fraction of tree per generation / sum of all fractions of tree per generation at the same time point). Each square represents the cells of a tree that are in the respective generation at the given time point. The redder the box is, the higher is the fraction of the tree in the respective generation. Thus, red boxes indicate synchronized cell divisions whereas green boxes indicate desynchronization. Samples from the continuous stimulation setting are depicted in the upper part. Below that are the short stimulation samples. The short stimulation + IL-12 samples are depicted in the bottom part. **(E)** As in (A) but αCD25-APC was added to the culture. The inter-division times of all cells from all investigated trees were separated according to their generation, and their inter-division times were plotted against their CD25 expression. Blue: brief stimulation without IL-12. Red: brief stimulation with IL-12. Green: continuous stimulation. Spearman r and p-values as indicated. Results from: Generation 1: 80 cells, generation 2: 138 cells, generation 3: 232 cells, generation 4: 313 cells, generation 5: 422 cells, generation 6: 583 cells. **(F)** Exemplary tree (brief stimulation + IL-12) shown as family tree (left panel. Loose ends with red X: cell died) and heat tree (right panel). Each box represents a cell as in the family tree. Yellow: low CD25 expression. Red: high CD25 expression. Blank: no CD25 quantification possible.

### Adoption of slower division speed coincides with expression of CMP marker CD62L

Finally, we aimed to investigate whether these linked changes in IL-2 receptivity and division speed also correlated with memory vs. effector T cell differentiation. Since CD62L^+^ CMPs have been identified as early as day four post immunization *in vivo* and have been shown to divide slower than their CD62L^-^ counterparts (8), we decided to investigate expression of this memory marker in our experimental system. First, we analyzed single-cell-derived T cell responses via flow cytometry at day five after *in vitro* activation. In line with previous observations made *in vivo*, we found that larger T cell families contained lower percentages of CD62L^+^ T cells **(Fig. 4 A)**. Moreover, when tracking T cell responses derived from populations of 100 T cells via intracellular dye dilution, we found that those T cells which had undertaken more divisions contained lower percentages of CD62L^+^ cells compared to their less-divided counterparts **(Fig. 4 B and C)**. When tracking single-cell-derived T cell responses in the same manner, we found that T cell families stretched out across multiple generations and that CD62L^-^ cells accumulated in the stronger divided offspring of the same starting cell **(Fig. 4 D and E)**. While this observation could indicate that CD62L^-^ T cells divide faster, it could also mean that differentiation into CD62L^-^ T cells happens only after a certain generation is reached. To resolve this question, we again turned to continuous imaging. Immediately after T cell activation CD62L is enzymatically removed from the surface of activated T cells (25). In line with this shedding of CD62L, we found virtually no surface expression of this molecule immediately after T cell activation. However, CD62L surface expression reappeared in subsequent generation and coincided with a slower division speed relative to CD62L^-^ T cells found in the same generation **(Fig. 4 F)**. Importantly, division speed of CD62L^+^ T cells remained relatively fixed throughout generations, arguing that the gradual slow-down of T cell families that we had observed from generations 3 to 6 resulted from an increasing number of T cells switching into the CD62L^+^ state. The same differentiation associated reduction of cell cycle speed occurred in presence of IL-12, albeit with CD62L^+^ T cells maintaining substantially faster division activity than in absence of this inflammatory cytokine **(Fig. 4 G)**.

**Figure 4:**
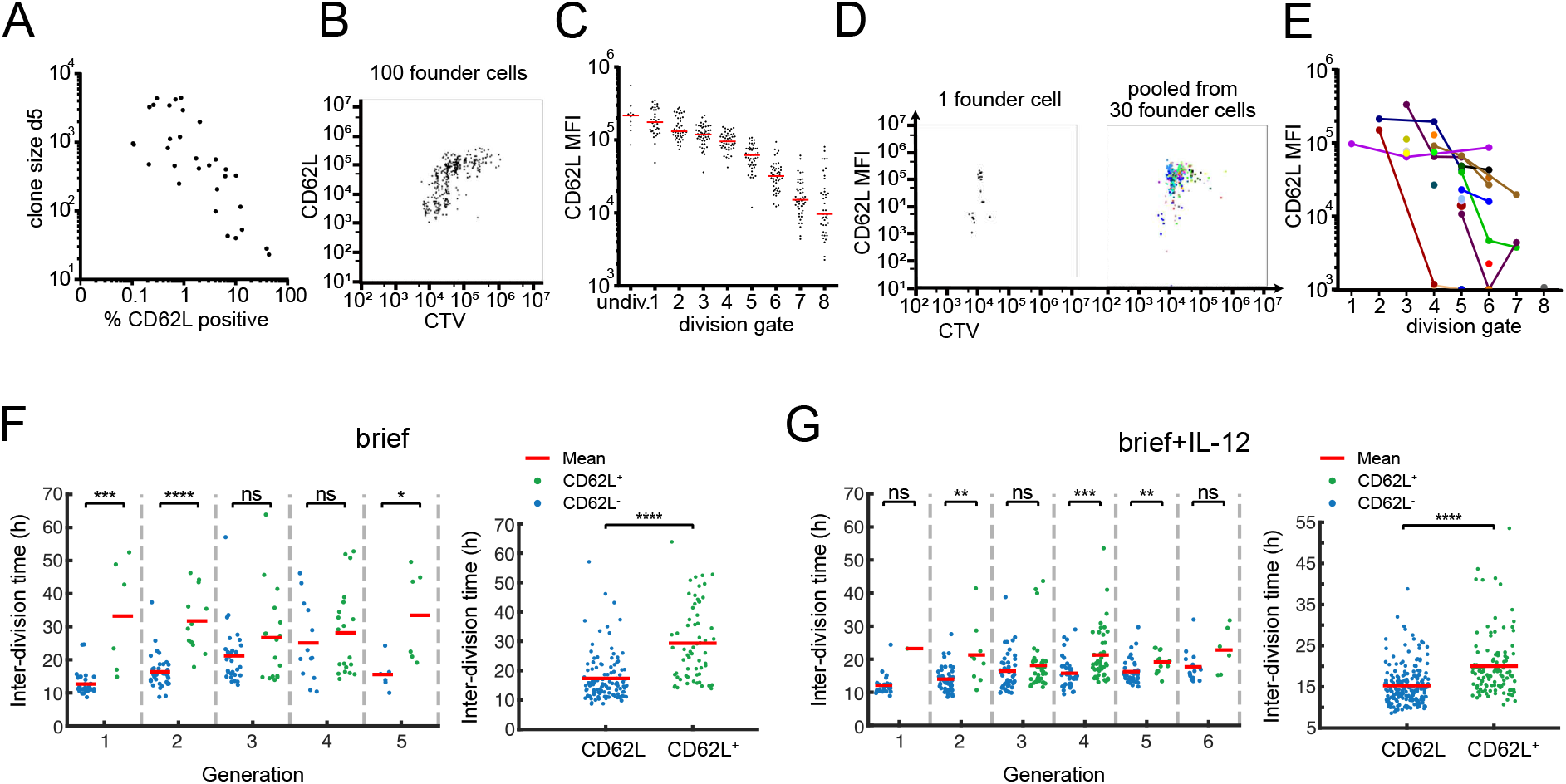
Adoption of slower division speed coincides with expression of CMP marker CD62L. **(A)** Cell numbers of single cell-derived progenies are plotted against the percentage of CD62L expressing cells (determined by flow cytometry) within the respective T-cell family (continuous TCR stimulation). Spearman correlation coefficient r = −0.7569, p < 0.0001 n = 31 (cells with %CD62L = 0% excluded from analysis). **(B)** Representative Cell Trace Violet plot of the offspring of 100 OT-I cells that had been stained with Cell Trace Violet and activated for 24 h. After further 3 days of cell culture, the cells were stained for CD62L and analyzed by flow cytometry. **(C)** On the basis of the vertical lines in the Cell Trace Violet plot in (B), the number of cell divisions for each cell in the plot can be estimated. The mean CD62L expression within each division peak is shown for all cells of the 100 cell-derived progenies. **(D)** Left panel: Representative Cell Trace Violet plot as in (B) but from a single progenitor cell. Right panel: Overlay of 30 single-cell Cell Trace Violet plots. Each color represents a different single-cell derived progeny. **(E)** CD62L expression in division peaks as in (C) but from single-cell derived progenies. In addition, division peaks of the same single-cell tree are drawn in the same color and connected with a line. **(F)** Inter-division times in different generations (left) and from all generations together (right) in the “brief” stimulation condition. The two-sided Wilcoxon rank sum test is used to determine if the inter-division times of CD62L^+^ (green) and CD62L^-^ (blue) cells are significantly different: ns: p-value > 0.05, *: p-value ≤ 0.05, **: p-value ≤ 0.01, ***: p-value ≤ 0.001, ****: p-value ≤ 0.0001. **(G)** Same as (F) for the “brief+IL-12” stimulation condition.

## Discussion

It is a hallmark of adaptive immunology that single antigen-specific T cells can generate progeny that diversifies into both terminally differentiated effector cells and precursors of long-lived memory cells (26). This fate diversification occurs in parallel to rapid T cell proliferation and computational modelling of single-cell-derived T cell responses has suggested that memory precursors and short-lied effector T cells are set apart by fundamentally distinct cell cycle speeds (2). Recently, we have shown that four days after initial T cell priming, memory precursors are characterized by a slower division speed than their effector counterparts *in vivo* (8). Early clonal expansion on the other hand was found *in vitro* to occur in a highly synchronized manner (15–18). Here, we set out to close the gap between these early *in vitro* observations and the later diversification of division speed and T cell fate observed *in vivo*. Therefore, we utilized continuous live-cell imaging and tracked the division speed, differentiation status and genealogical connections of all descendants derived from a single naïve T cell for up to ten divisions of activation-induced proliferation *in vitro*. We find that initial T cell proliferation indeed occurs in a burst-like manner. With rapid execution of T cell divisions being largely independent of further TCR and cytokine stimuli received beyond priming. At two to three cell divisions, the average duration of this burst is similar to that previously described for *in vitro* settings in which stimuli were stringently restricted to the starting cell (18, 27, 28). Upon restriction of subsequent stimuli, including blockade of endogenous IL-2, T cell proliferation abruptly subsides after completion of this burst (18, 27, 28). This homogenous programming of proliferation activity, termed “division destiny”, has been attributed to a division counter (18, 27) or division timer (28) set in the starting cell and transmitted to all of its progeny. Here, we deliberately chose culture conditions that maintained T cell proliferation beyond the initial burst phase and closely monitored the evolution of cell cycle speed within the resulting family trees. Interestingly, we found evidence for a heredity of cell cycle speed that was in part programmed in the starting cell but also critically required the emergence of distinct T cell subsets proliferating at distinct speeds. One of these, the EA subset, was of transient nature and its life-time coincided with the duration of the initial proliferative burst. Thereafter, slow (S) and fast (F) cycling subsets emerged that were characterized by distinct CD25 and CD62L expression levels, two markers that have been found to characterize the early subdivision of long-lived CMPs (CD62L^+^CD25^-^) and non-CMPs (CD62L^-^CD25^+^) *in vivo* (29, 30). In keeping with a concept of asymmetric cell division (31), the segregation of these subsets has been proposed to occur as early as the first cell division. While our *in vitro* data cannot exclude such an immediate segregation, they rather support a developmental model in which lineage segregation begins somewhat later, after a burst-like expansion of two to three cells division. Thereafter, we find that individual branches within an expanding T cell family tree can maintain slower or faster cell cycle speeds that coincide with differences in expression of CD25 and CD62L. With these findings, our study puts renewed focus on the intertwined nature of cell cycle activity and T cell differentiation (32, 33). While previously this relation has been mainly explored with respect to the accumulated number of divisions, we now highlight the actual speed of cell division as a major heritable property that appears to be regulated in parallel to key lineage decisions of activated T cells. From a methodological point of view, we provide a novel computational inference framework for analyzing tree-structured data obtained by live-cell imaging. Due to the complexity of such tree-structured data, previous studies have mostly utilized summary statistics from live-cell imaging experiments to infer underlying kinetics (15, 34, 35). Our framework, however, exploits full structural information from this data type. As opposed to few other model-based analyses of lineage trees, where phenotypic measurements were assumed to inform about cellular differentiation (36–39), our framework does not rely on phenotypic observations and instead links division speed to underlying hidden states. Unlike similar studies (38), an important feature of our computational modeling is the simultaneous analysis of complete genealogical trees without partitioning into smaller “sub-trees”; this ensures that no long-ranged structural information is lost. Our inference framework allows to investigate the complex kinetic structure of expanding T cell family trees and enables hypothesis testing as to the hereditary nature of cell cycle activity and the topological organization of T cell differentiation. It will be exciting to investigate whether the developmental framework proposed here will hold true *in vivo* and to more closely examine how T cell differentiation and cell cycle speed are connected on the molecular level.

## Acknowledgements

This work was supported by the ‘European Research Council starting grant’ (949719 – SCIMAP) to V.R.B., the German research foundation (DFG) – SFB 1054 (Project number 210592381) to V.R.B. and D.H.B. and the BMBF project Quan-T-cell (e:Med initiative on Systems Medicine, FKZ 01ZX1505) to M.F.

## Author contributions

V.R.B., M.F. and D.H.B. designed and supervised the study. V.R.B., A.K. and M.P. wrote the manuscript. M.P. and D.L. conducted experiments. A.K. and M.F. performed computational analyses and developed the bayesian framework for analysis of tree structured data. MP and D.L. performed continuous imaging. T.S. supervised continuous imaging and contributed to the study design.

## Declaration of competing interests

The authors declare no competing interests.

## Material and Methods

### Mice

C57BL/6 wt mice were purchased from Envigo. OT-I Rag1-/- matrix donor mice expressing combinations of the congenic markers CD45.1/2 and CD90.1/2 were bred under specific pathogen-free conditions at the mouse facility of the Institute for Medical Microbiology, Immunology and Hygiene, Technical University of Munich (TUM), Munich 81675, Germany. Animal care and procedures were in accordance with institutional protocols as approved by the relevant local authorities.

### Cell Sorting

Cells were isolated from peripheral blood or spleens of C57BL/6 or OT-I Rag1-/- matrix donor mice and sorted for a naive phenotype (CD8+, CD44low) at a MoFlo Legacy or MoFLo XDP cell sorter. For experiments with sustained anti-CD3/CD28 stimulation a single naive CD8+ T cell was sorted per well of an anti-CD3/CD28 coated (10 μg/mL at 4°C over night) plate (384 Well Small Volume™ LoBase med. binding μClear^®^ microplate) containing 25 μL RPMI + Pen/Strep + 10 % heat-inactivated FCS and 25 U/mL recombinant human IL-2. For experiments with limited anti-CD3/CD28 stimulation 10 000 naive CD8+, CD44low T cells were sorted per well of an anti-CD3/CD28 coated (10 μg/mL at 4°C over night) plate (384 Well med. binding μClear^®^ microplate) containing 100 μL RPMI + Pen/Strep + 10 % heat-inactivated FCS + 25 U/mL recombinant human IL-2 + 10 ng/mL murine IL-12. After 24 h at 37°C + 5% CO2 + 95% H2O the cells were pooled and sorted again. This time it was sorted for activated (CD8+, CD44high) T cells and a single activated cell was sorted per well of an anti-CD28 coated 384 well plate (384 Well Small Volume™ LoBase med. binding μClear^®^ microplate) containing 25 μL RPMI + Pen/Strep + 10 % heat-inactivated FCS and 25 U/mL recombinant human IL-2 with or without 10 ng/mL murine IL-12.

### Continuous Single-Cell Imaging

Live-cell imaging was performed by the ETH Zurich as described by Eilken et al. (20). Briefly, the microplates were imaged using a Nikon-Ti Eclipse with linear encoded motorized stage, Orca Flash 4.0 V2 (Hamamatsu), and Spectra X fluorescent light source (Lumencor) and 10x CFI Plan Apochromat λ objective (NA 0.45). White light emitted by Spectra X was collimated and used as a transmitted light for bright field illumination via a custom-made motorized mirror controlled by Arduino UNO Rev3 (Arduino) and images were taken every 1-3 minutes. For experiments in which the expression of a surface antigen was measured, the respective dye-conjugated antibody was added in very low concentration to the culture medium (e.g., 1:20 000 for anti-CD25-APC, corresponding to 10 ng/mL) and in addition to the bright field images the respective fluorescent channel was acquired approximately once every 45 minutes using filtersets for PE (546/10; 560LP; 577/25) and APC (620/60; 660LP; 700/75; both AHF). The cells were observed for 3-5 days at 37°C, 5% CO2 and 5% O2. All images were acquired a 16-bit lossless .png and linearly transformed using optimal black

### Cell Tracking

Generation of T cell family trees and heat trees was performed by manually tracking the cells using “The Tracking Tool” and the semi-automatic FI-measurement software “qTFy” (Hilsenbeck, O., et al., Software tools for single-cell tracking and quantification of cellular and molecular properties. Nat Biotechnol, 2016. 34(7): p. 703-6.)

### Cell Proliferation Dye Staining and Flow Cytometry

Cell Proliferation Dye staining was performed using the CellTrace Violet™ Cell Proliferation Kit by ThermoFisher scientific according to the manufacturer’s instructions. Cells were sorted, activated for 24 h and sorted again as described in the section about cell sorting. After three days the cells were stained with anti-CD62L-FITC (MEL-14) and acquired using a Cytoflex S cytometer.

### ELISA (Suppl.)

The IL-2 ELISA was performed using the IL-2 Human ELISA Kit by ThermoFisher scientific according to the manufacturer’s instructions.

### Antibodies and Cytokines

Anti-murine CD3 (145-2C11), anti-murine CD28 (37.51) and anti-murine CD25-APC (PC61) were purchased from BD Bioscience, anti-murine CD8 (53.6.7), anti-murine CD44 (IM7) and anti-murine CD62L-FITC (MEL-14) were purchased from Biolegend. Recombinant murine IL-12 was purchased from Peprotech. Recombinant human IL-2 was purchased from ThermoFisher Scientific.

### Statistical Analysis

All non-in silico experiments were analyzed using Prism 7, 8 and 9 (GraphPad Software). p-values were assessed using a two-tailed unpaired Student’s t test, one-way ANOVA, Spearman non-parametric or Pearson test, as specified in the figure legends.

---

**Figure S1:**
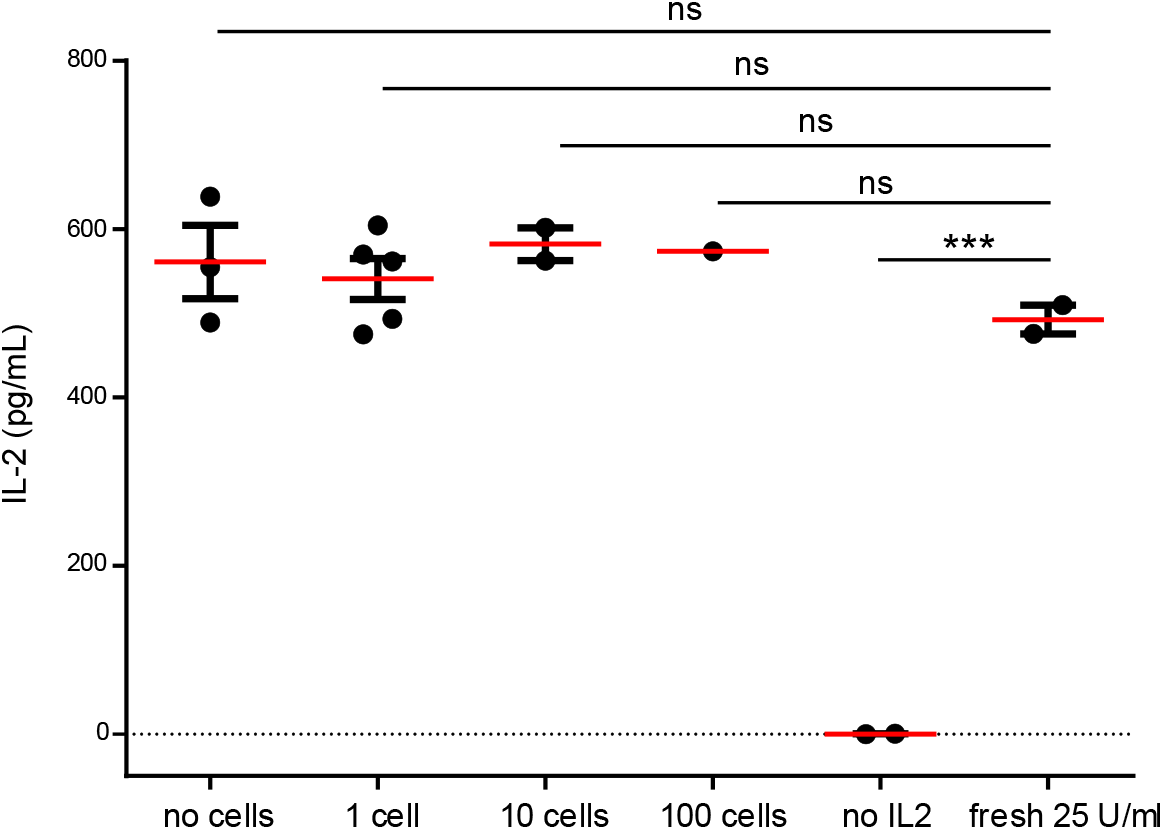
IL-2 levels are stable throughout the experiment. Cells were activated for 24 h. Subsequently, 0, 1, 10 or 100 cells were transferred into wells with 25 U/mL IL-2 and cultured for further three days. IL-2 in the supernatant was detected by ELISA. As a control, culture medium without cells was measured, containing no or 25 U/mL freshly added IL-2. One-way ANOVA and Dunnett’s multiple comparison test: **** p<0.0001.

**Figure S2:**
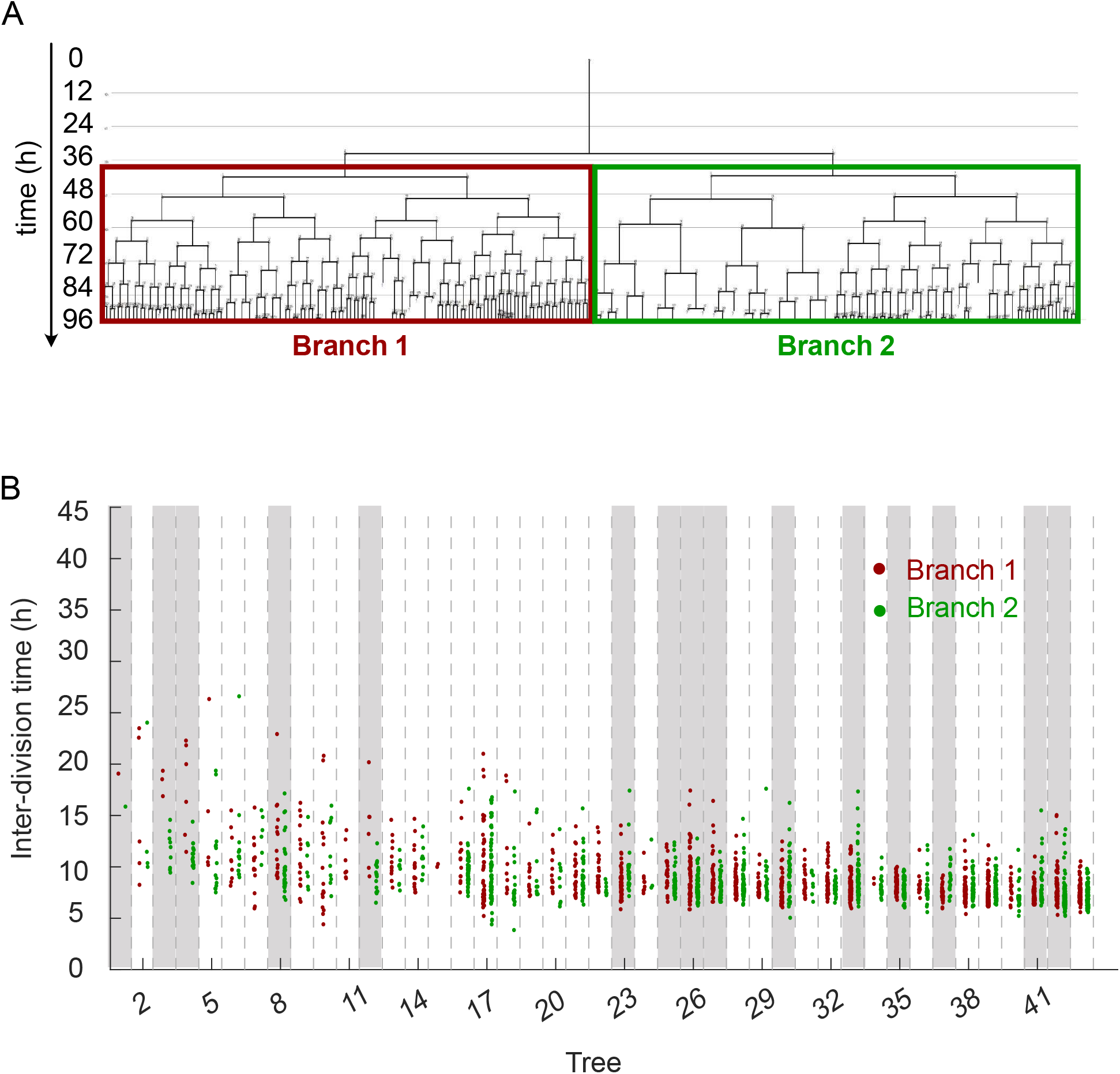
Comparison of division speeds in the two branches emerging after the first cell division. **(A)** Analogue to Figure 1I: A representative family tree was divided into two branches (red and green) after the first cell division. **(B)** Inter-division time of cells in the two branches starting from the first generation are color-coded for all trees in Figure 1F. In 15 out of 43 trees (~35%, tress highlighted in gray), interdivision times differed significantly between the two branches. For every tree, a p-value was calculated based on one-way ANOVA. We then estimated positive false discovery rates (pFDR) for multiple hypothesis testing based on these p-values, and used a cutoff of pFDR<0.05 for significance.

**Figure S3:**
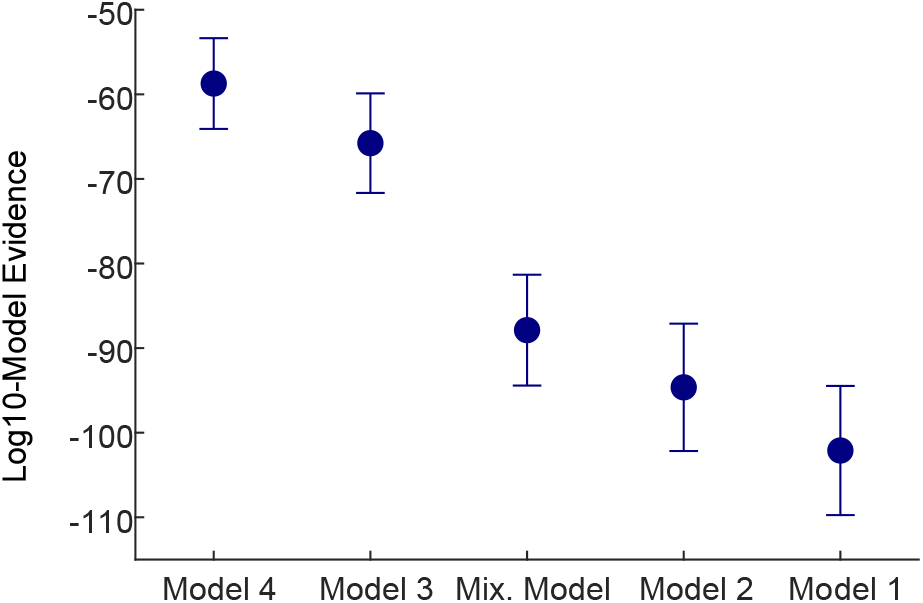
Model comparison summary from the results of all data groups. The model evidences for models #1, #2, #3, #4 (Fig. 1 C and J) and the mixture model indicate the following hierarchy where model #4 explains the data best and model #1 is least matching to the data: model #4 > model #3 > mixture model > model #2 > model #1. The circles show the mean of model evidences of eight data groups (Supplementary Methods). The error bars show the standard error of the mean.

**Figure S4:**
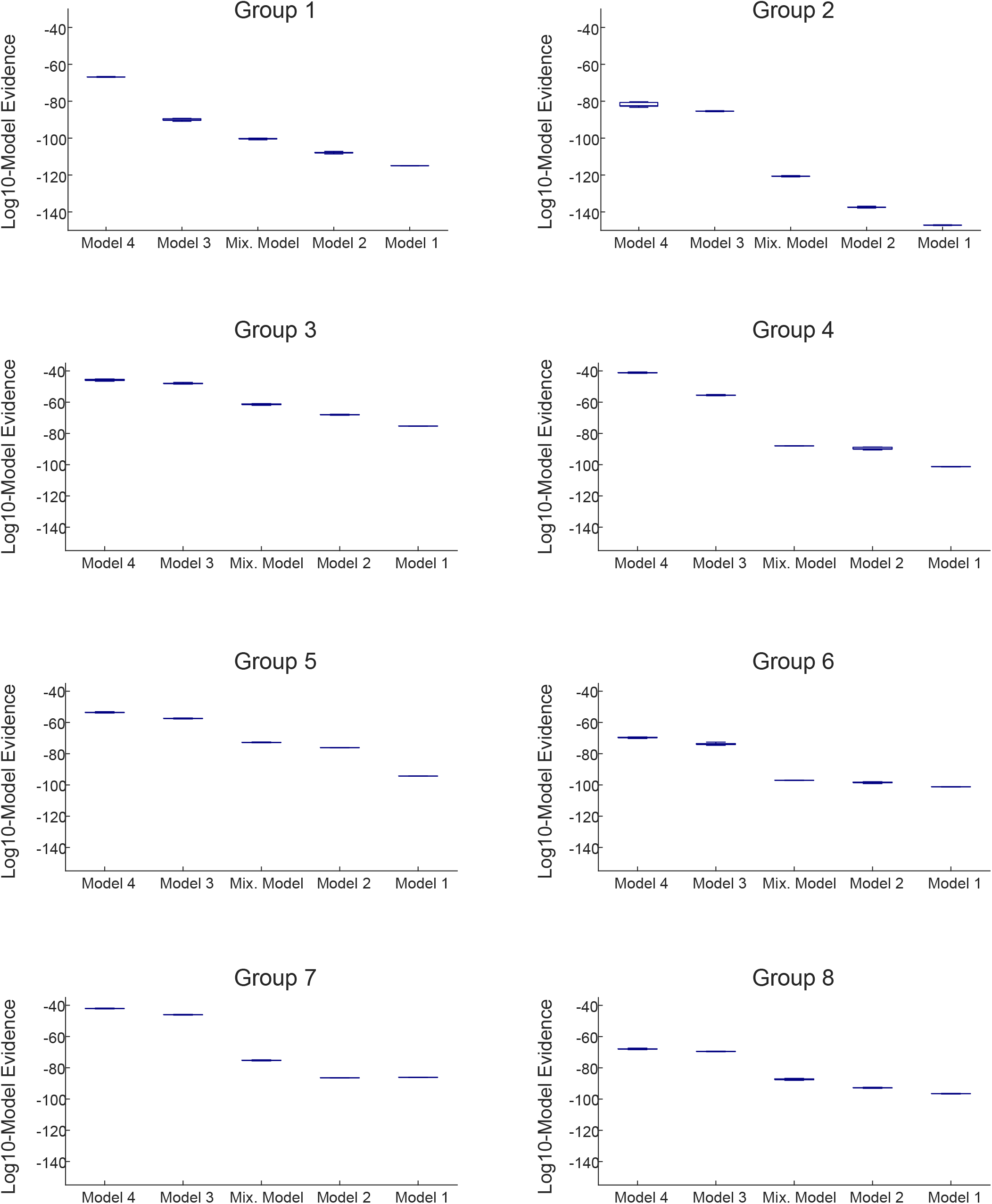
Model comparison results in individual groups. The model evidences for models #1, #2, #3, #4 (Fig. 1 C and J) and the mixture model for the eight data groups. The boxplots show the distribution of the model evidences calculated based on 10000 bootstrapped samples of parameter posteriors (Supplementary Methods).

**Figure S5:**
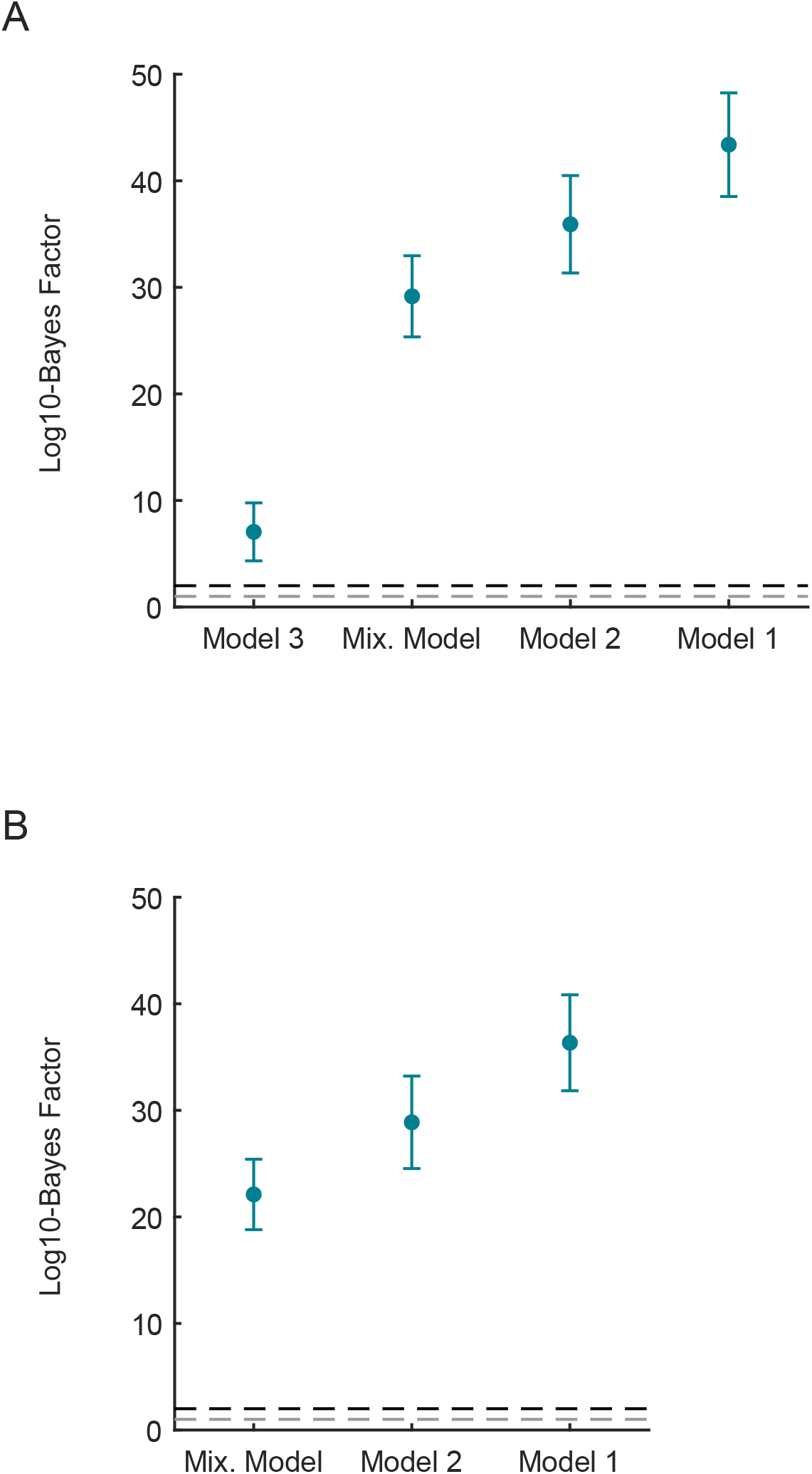
Bayes factors summary from the results of all data groups. **(A)** The log10-Bayes factor of model #4 (Fig. 1J) compared to models #1, #2, #3 (Fig. 1C) and the mixture model (Supplementary Methods). The circles show the mean of log10-Bayes factors calculated in the eight data groups. The error bars show the standard error of the mean. The dashed grey line shows the cutoff value for “strong evidence” (log10-Bayes factor = 1) and the dashed black line shows the cutoff value for “decisive evidence” (log10-Bayes factor = 2) for model #4. **(B)** Same as (A) where the log10-Bayes factors are calculated for model #3 compared to models #2 and #1 and the mixture model.

**Figure S6:**
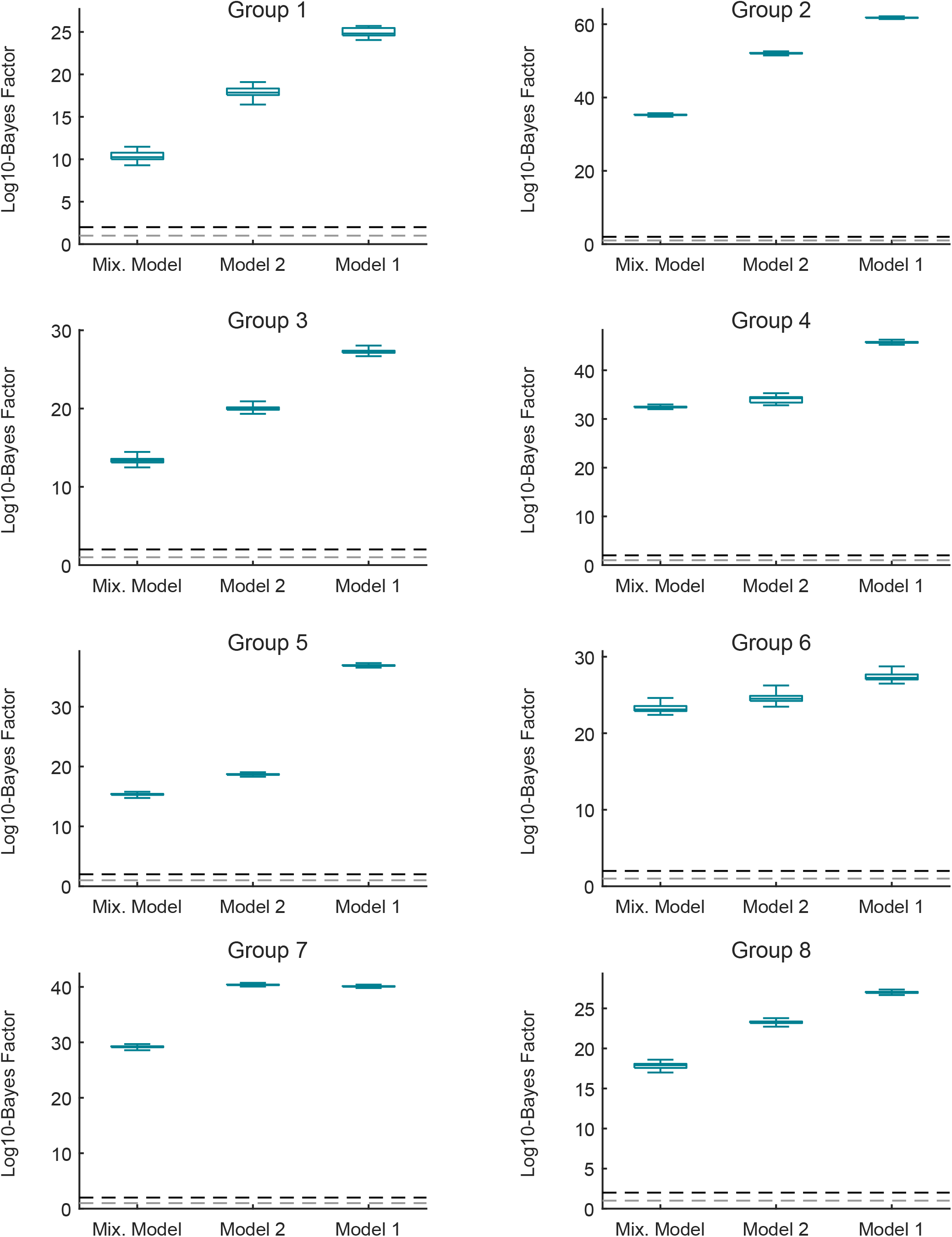
Bayes factors in individual groups with respect to model #3. The log10-Bayes factor of model #3 (Fig. 1C) compared to models #1, #2 (Fig. 1C) and the mixture model for the eight data groups. The boxplots show the distribution of the log10-Bayes factors calculated based on 10000 bootstrapped samples of parameter posteriors (Supplementary Methods). The dashed grey line shows the cutoff value for “strong evidence” (log10-Bayes factor = 1) and the dashed black line shows the cutoff value for “decisive evidence” (log10-Bayes factor = 2) for model #3.

**Figure S7:**
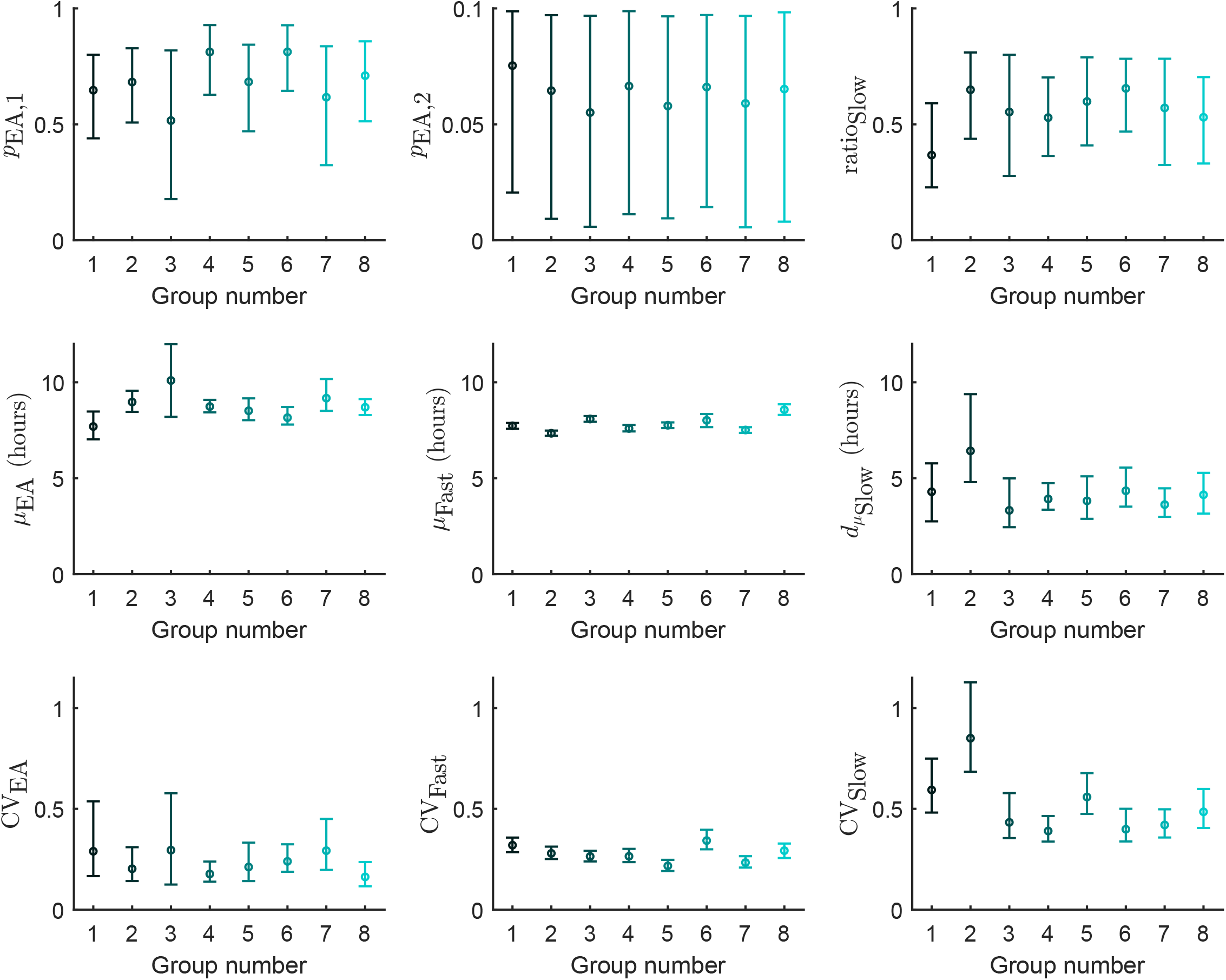
Inferred parameter values for model #3 in the eight data groups. The circles and the error bars respectively show the median and the 95% credible intervals of the parameter posterior distributions.

**Figure S8:**
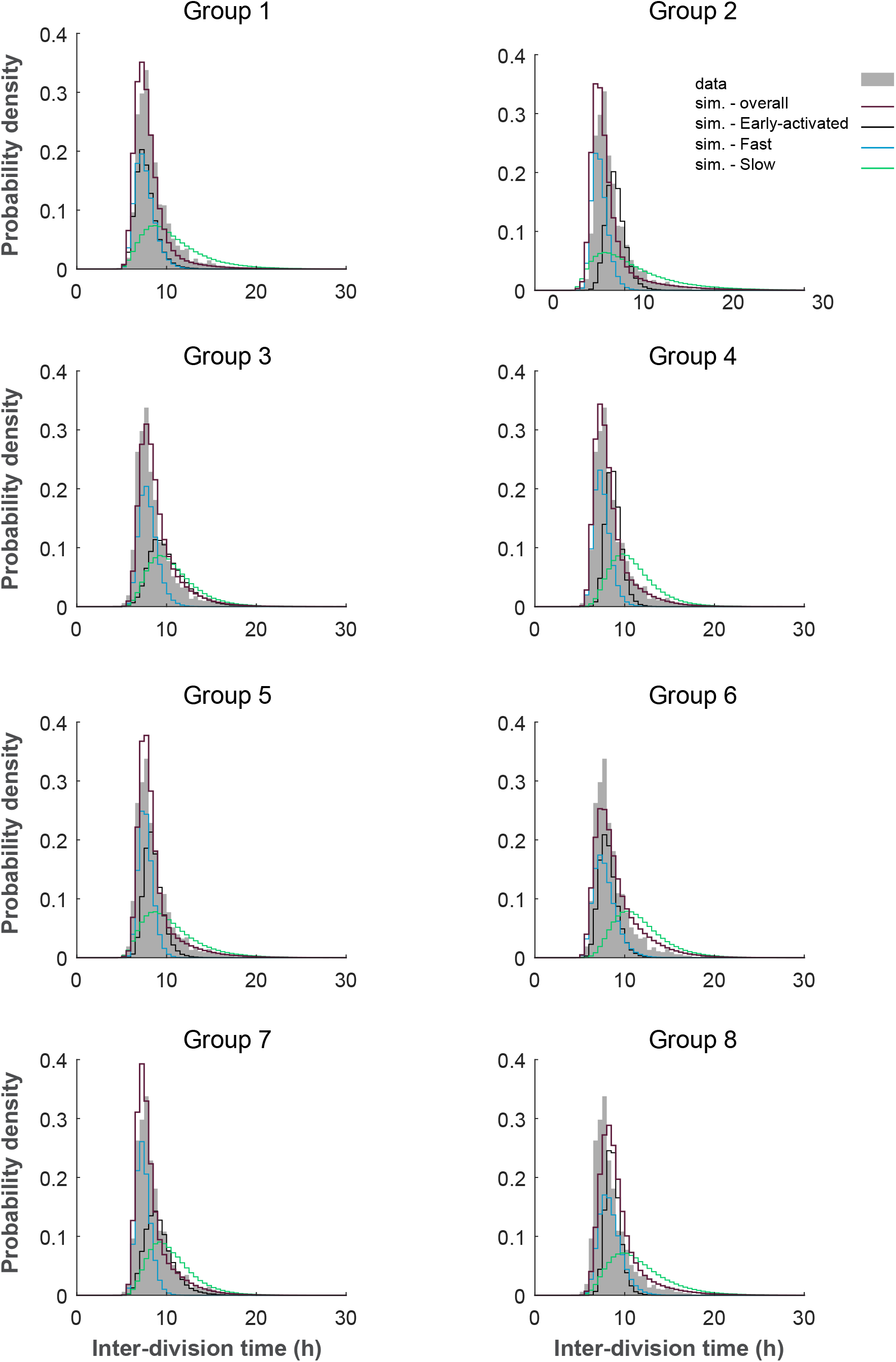
The distribution of inter-division times in simulated data based on model #3. The distribution of inter-division times in 10000 simulated trees based on model #3 compared to that of the experimental data (grey) for the eight data groups. In every group, the parameter posteriors inferred in that group are used for simulating the trees. The red histogram shows the overall distribution in the simulated data, while the black, blue and green histograms show the distribution of “Early-activated”, “Fast-dividing” and “Slow-dividing” subsets respectively.

**Figure S9:**
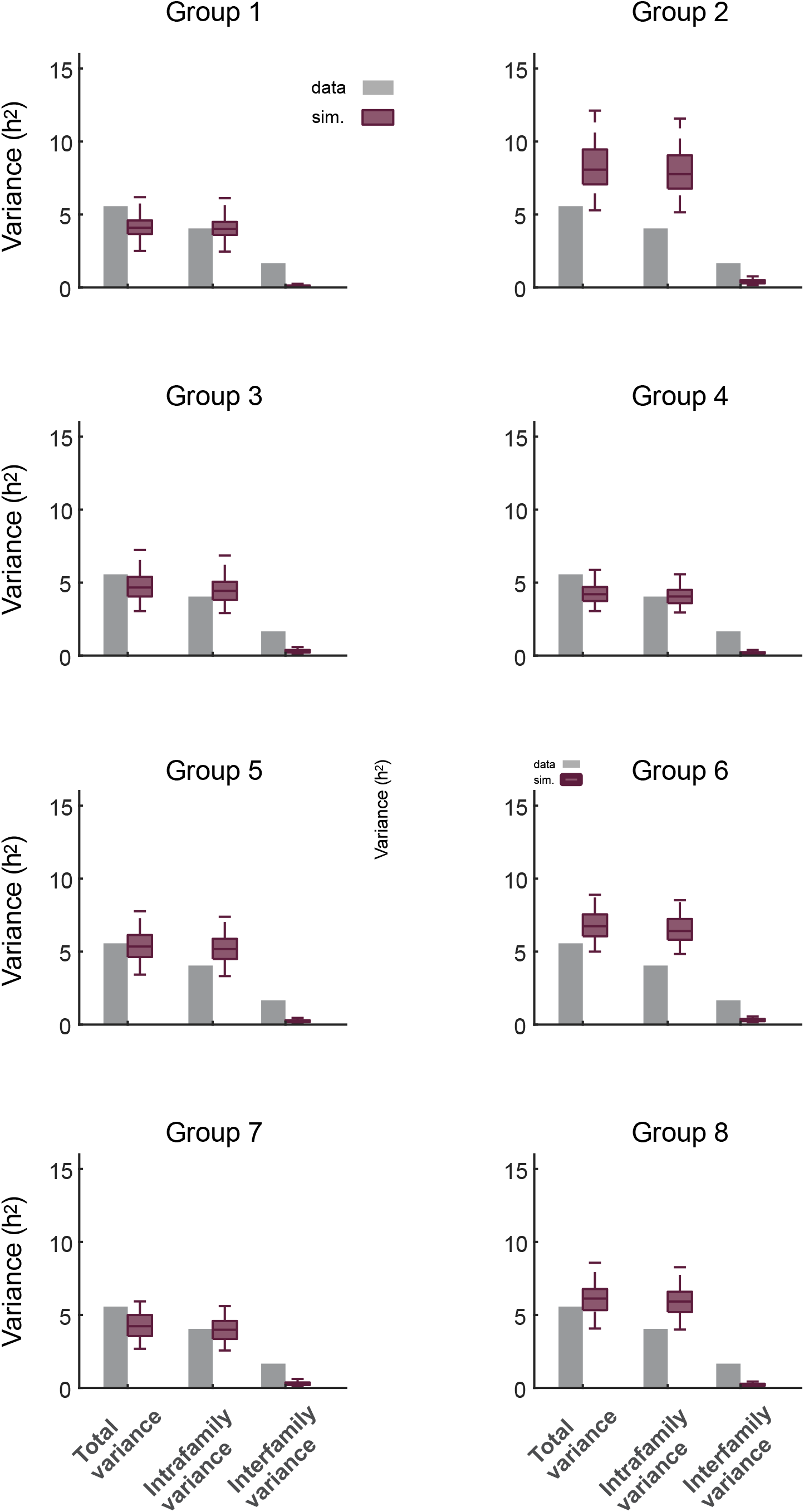
Variance of inter-division times in the simulated data based on model #3. Total variance of the interdivision times and the contribution of intrafamily and interfamily sources as observed in the experimental data (grey bars) and the simulated data (red boxes) for the eight data groups. The simulated data consists of 500 datasets of each 44 trees simulated based on model #3 and the parameter posteriors of every data group. Intrafamily variance is calculated as the weighted mean of the variances of interdivision times within different families. Interfamily variance is calculated as the weighted variance of the family mean inter-division times (Supplementary Methods).

**Figure S10:**
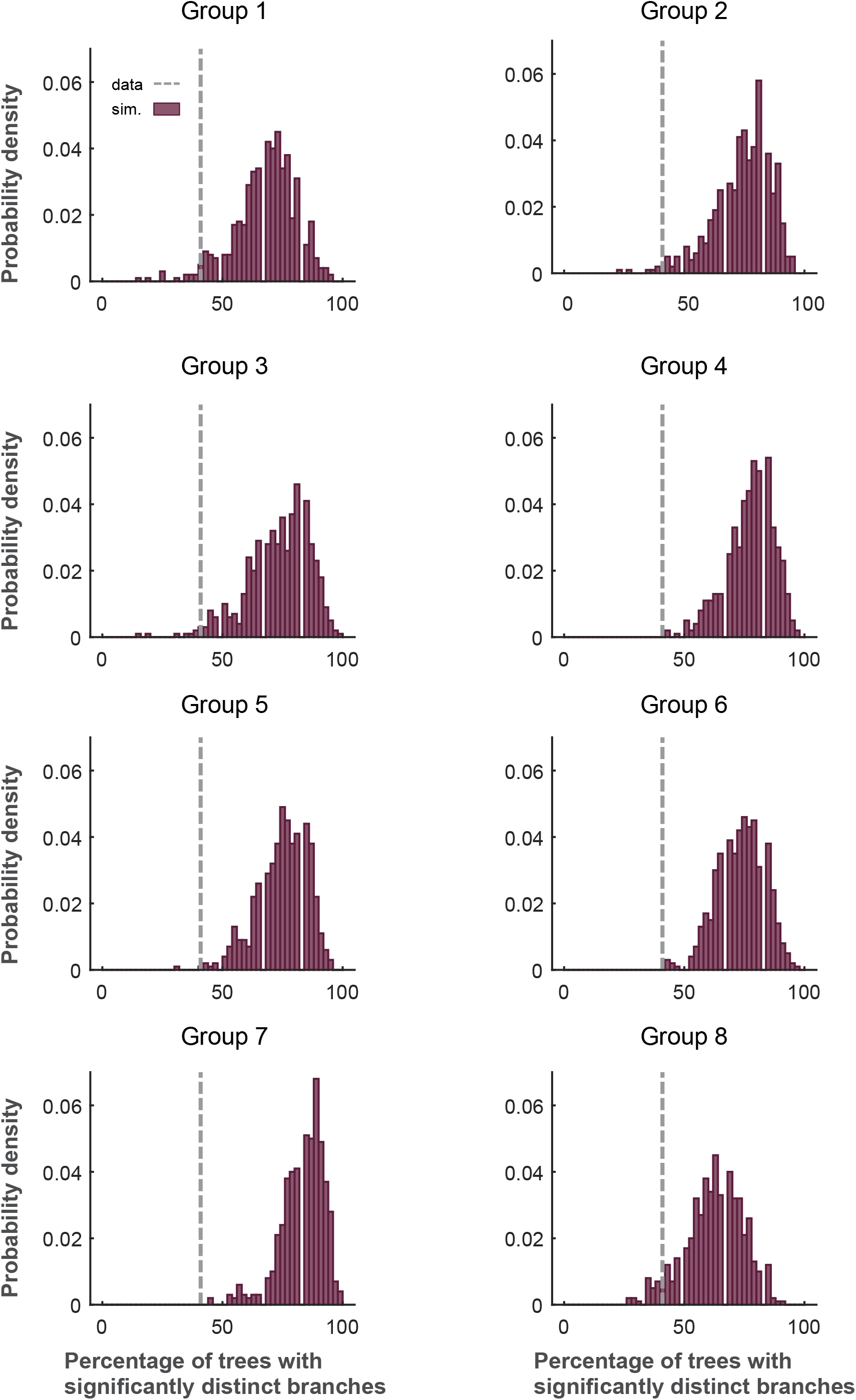
Percentage of trees with significantly distinct branches in the simulated data based on model #3. Percentage of the trees whose four branches (as in Fig. 1I) have significantly distinct inter-division times is shown. The grey line shows the experimental data and the red histogram shows the distribution of this percentage in the simulated data for the eight data groups. The simulated data consists of 500 datasets of each 44 trees simulated based on model #3 and the parameter posteriors of every data group. (Supplementary Methods).

**Figure S11:**
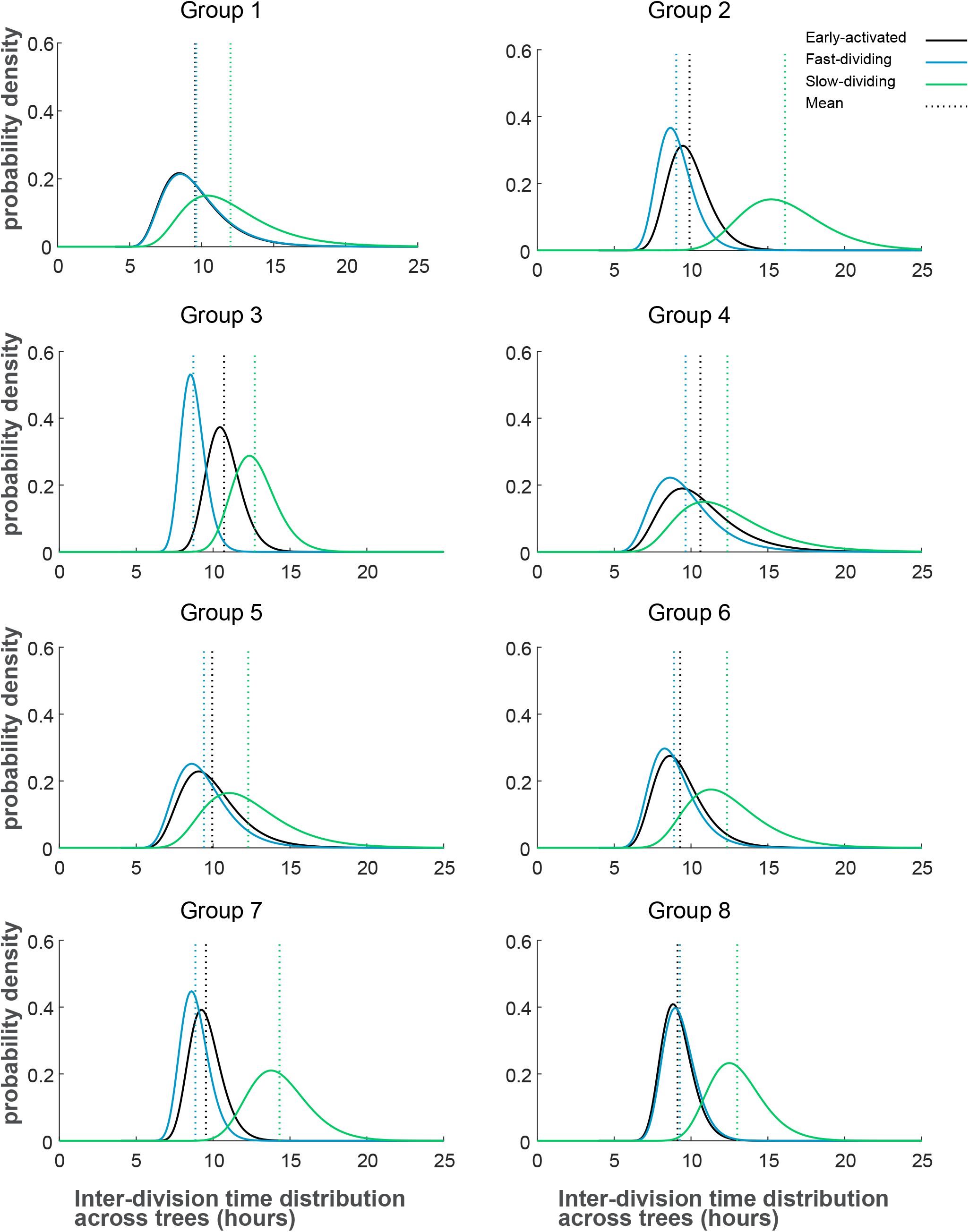
Variation of subset-specific mean inter-division times between different families. Inferred distribution of subset-specific mean inter-division times based on model #4 for the eight data groups. The black, blue and green curves show the distribution for the “Early-activated”, “Fast-dividing” and “Slow-dividing” subsets respectively.

**Figure S12:**
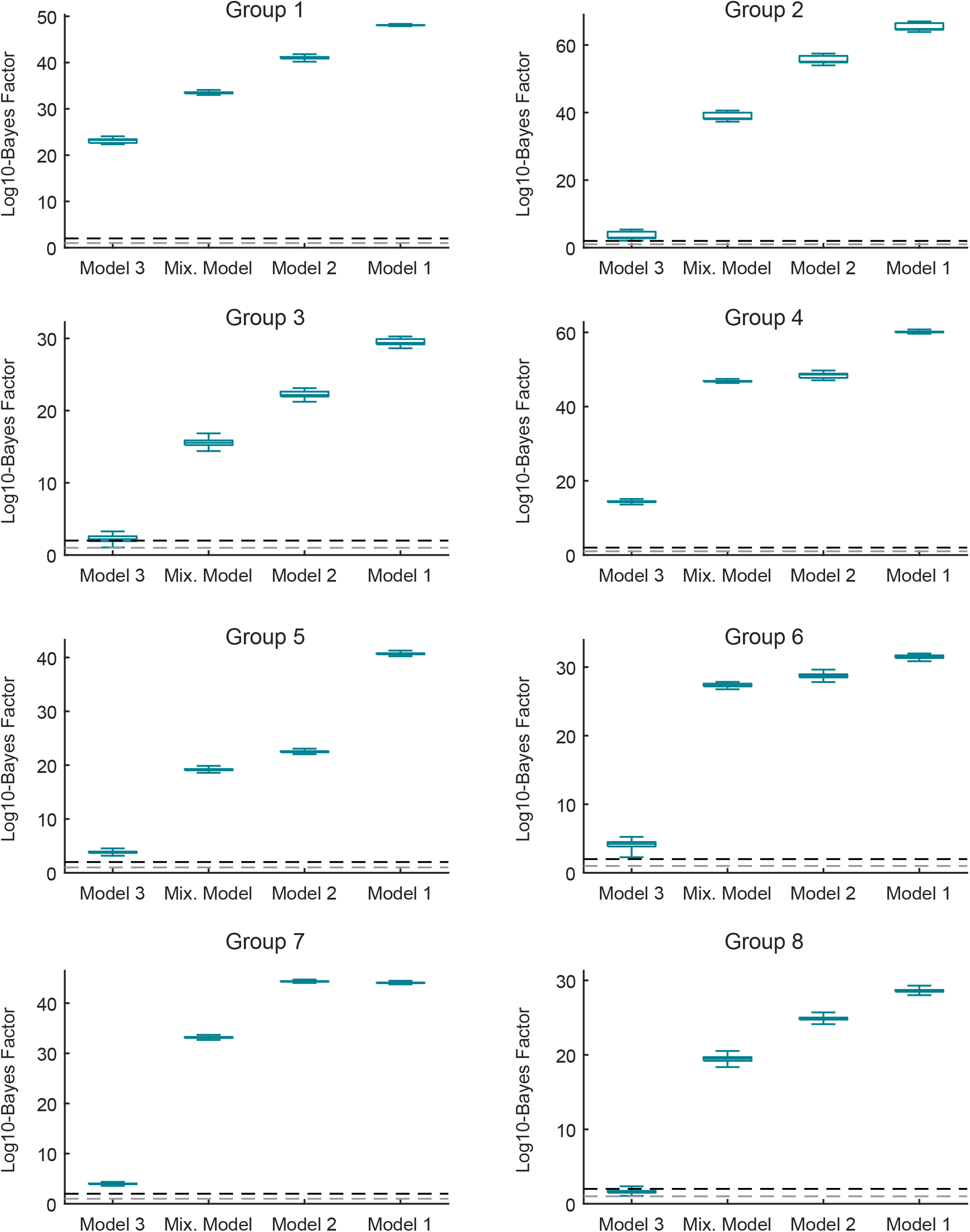
Bayes factors in individual groups with respect to model #4. The log10-Bayes factor of model #4 (Fig. 1C) compared to models #1, #2, #3 (Fig. 1C) and the mixture model (Supplementary Methods) for the eight data groups. The boxplots show the distribution of the log10-Bayes factors calculated based on 10000 bootstrapped samples of parameter posteriors (Supplementary Methods). The dashed grey line shows the cutoff value for “strong evidence” (log10-Bayes factor = 1) and the dashed black line shows the cutoff value for “decisive evidence” (log10-Bayes factor = 2) for model #4.

**Figure S13:**
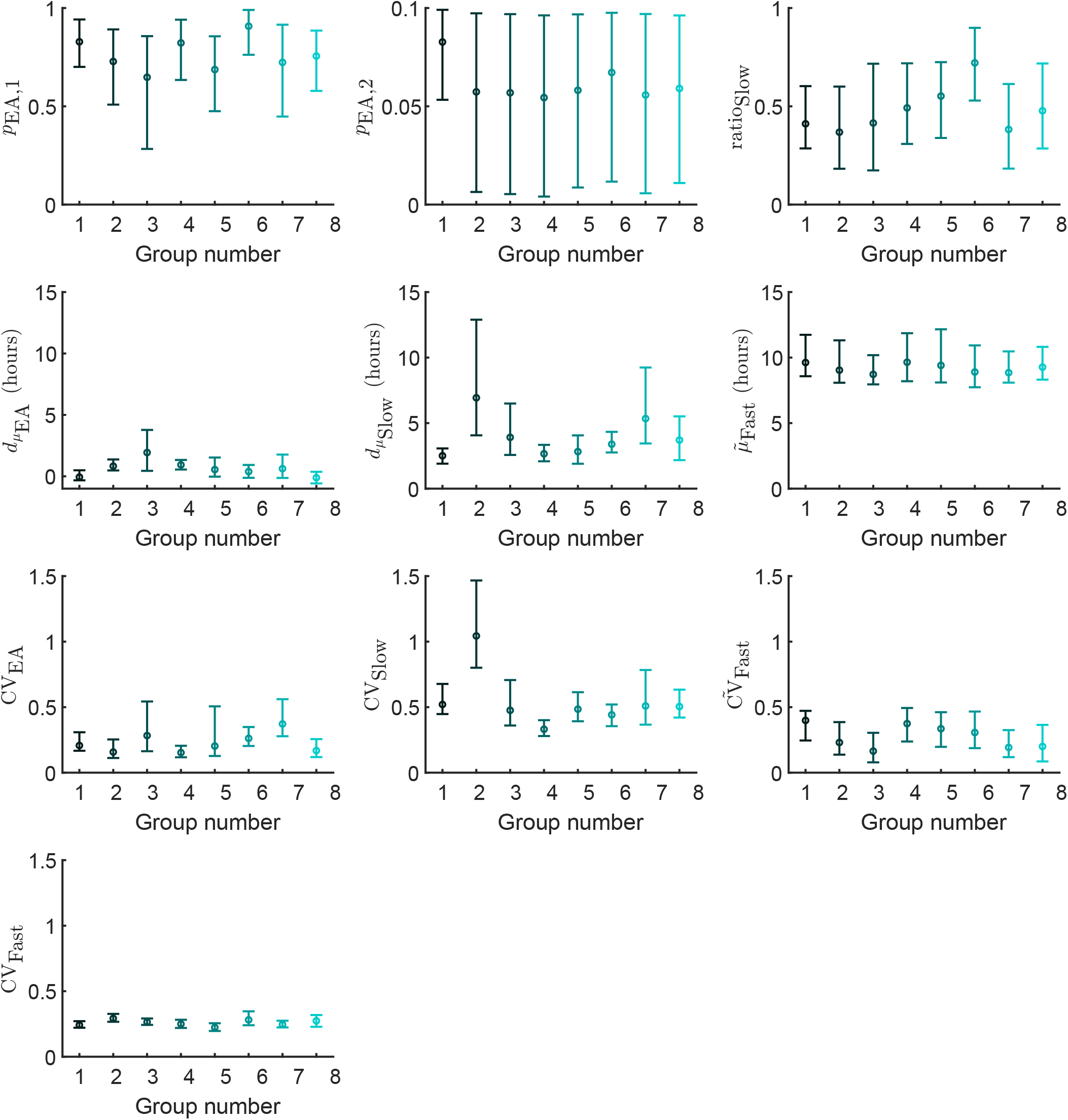
Inferred parameter values for model #4 in the eight data groups. The circles and the error bars respectively show the median and the 95% credible intervals of the parameter posterior distributions.

**Figure S14:**
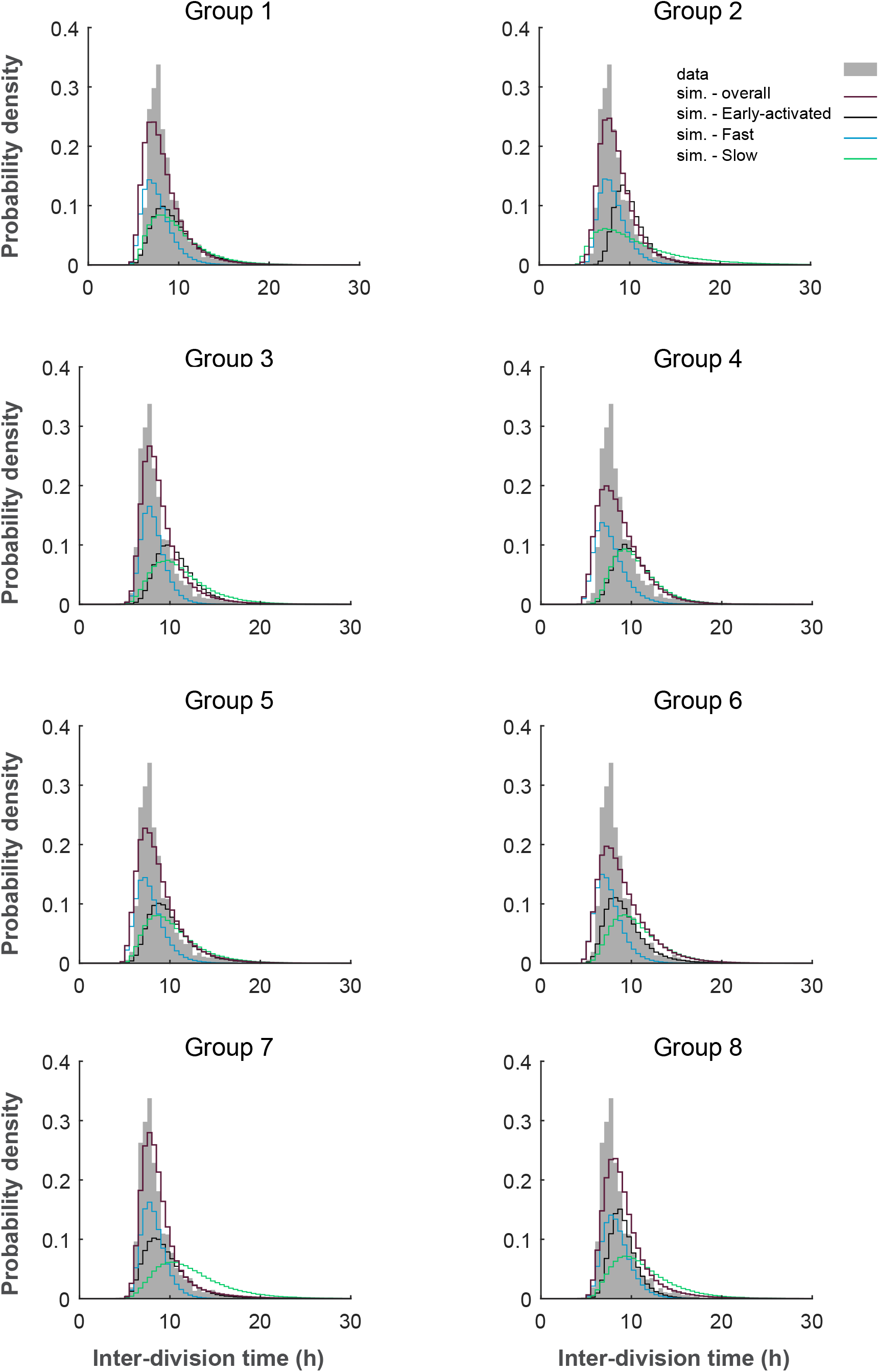
The distribution of inter-division times in simulated data based on model #4. The distribution of inter-division times in 10000 simulated trees based on model #4 compared to that of the experimental data (grey) for the eight data groups. In every group, the parameter posteriors inferred in that group are used for simulating the trees. The red histogram shows the overall distribution in the simulated data, while the black, blue and green histograms show the distribution of “Early-activated”, “Fast-dividing” and “Slow-dividing” subsets respectively.

**Figure S15:**
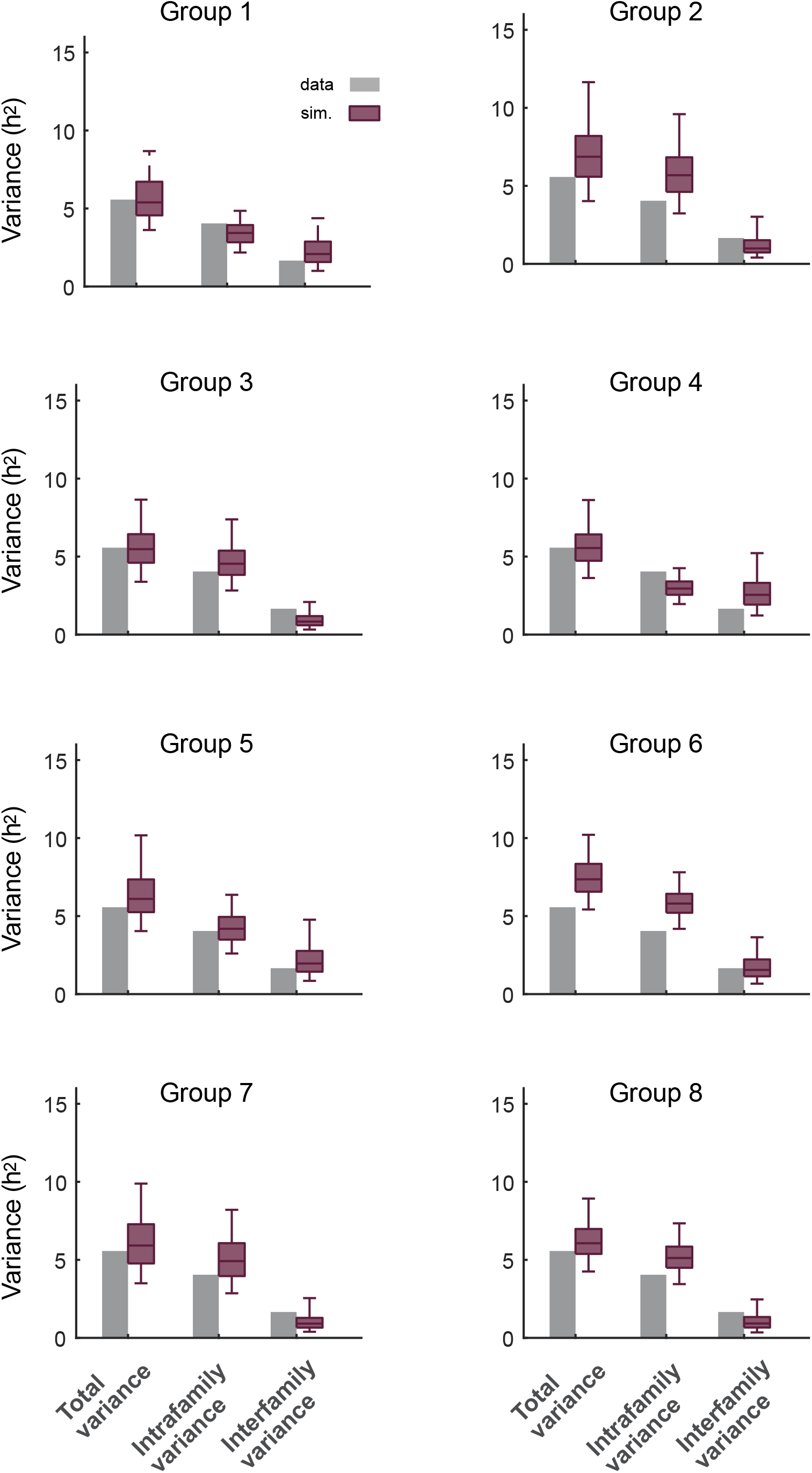
Variance of inter-division times in the simulated data based on model #4. Total variance of the interdivision times and the contribution of intrafamily and interfamily sources as observed in the experimental data (grey bars) and the simulated data (red boxes) for the eight data groups. The simulated data consists of 500 datasets of each 44 trees simulated based on model #4 and the parameter posteriors of every data group. Intrafamily variance is calculated as the weighted mean of the variances of inter-division times within different families. Interfamily variance is calculated as the weighted variance of the family mean inter-division times (Supplementary Methods).

**Figure S16:**
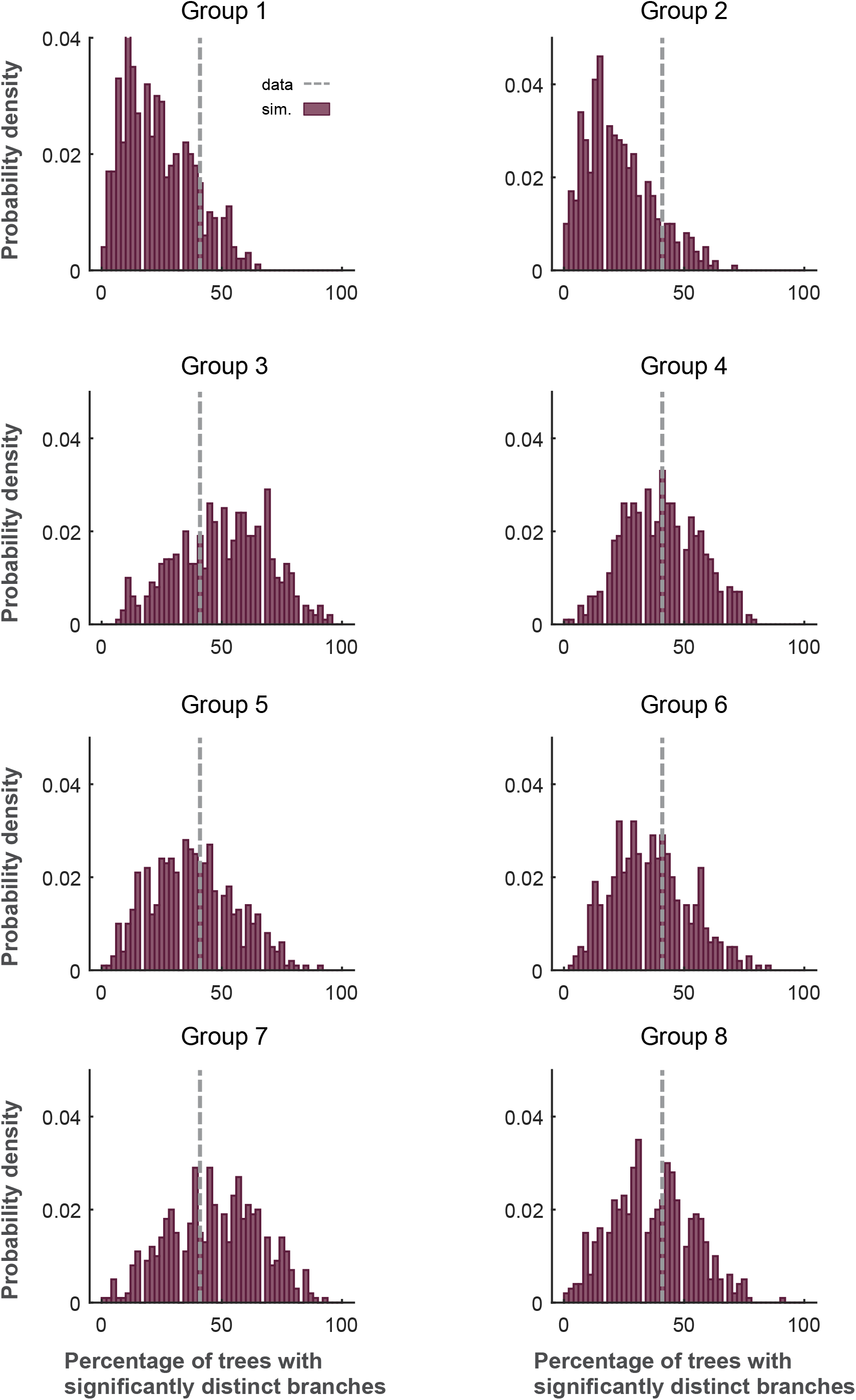
Percentage of trees with significantly distinct branches in the simulated data based on model #4. Percentage of the trees whose four branches (as in Fig. 1I) have significantly distinct inter-division times is shown. The grey line shows the experimental data and the red histogram shows the distribution of this percentage in the simulated data for the eight data groups. The simulated data consists of 500 datasets of each 44 trees simulated based on model #4 and the parameter posteriors of every data group. (Supplementary Methods).

**Figure S17:**
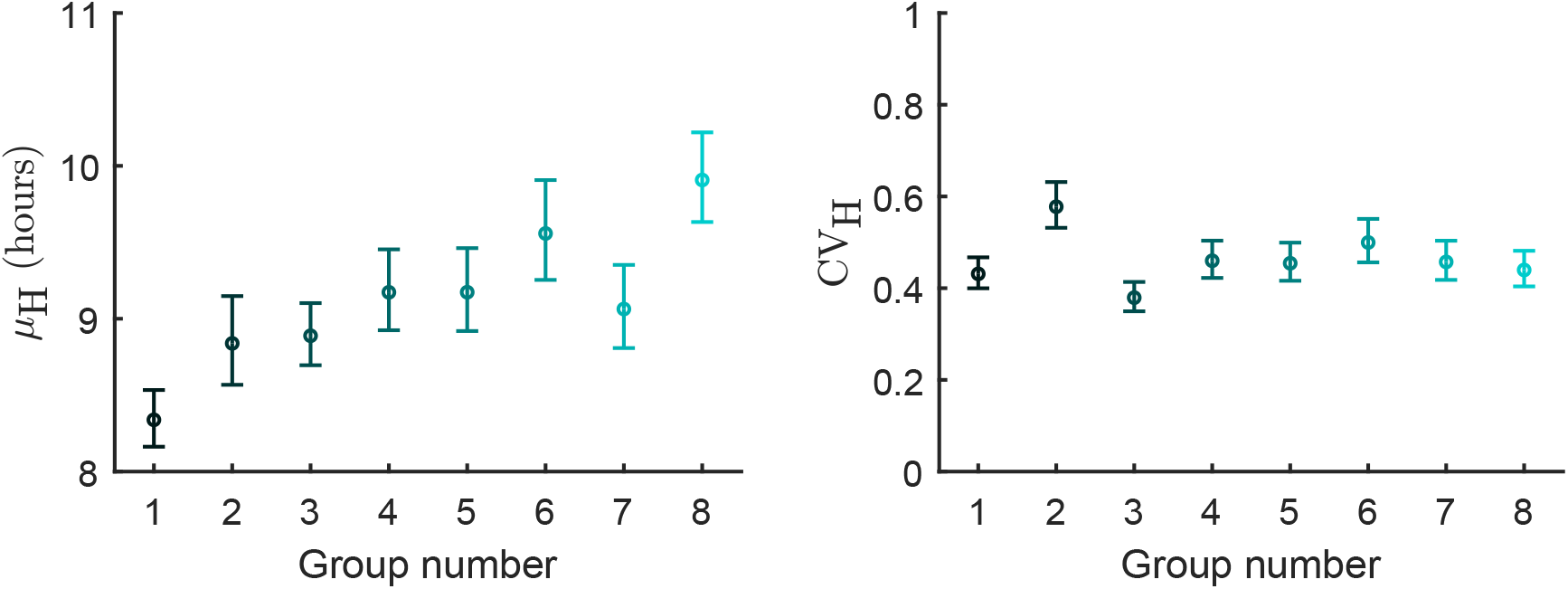
Inferred parameter values for model #1 in the eight data groups. The circles and the error bars respectively show the median and the 95% credible intervals of the parameter posterior distributions.

**Figure S18:**
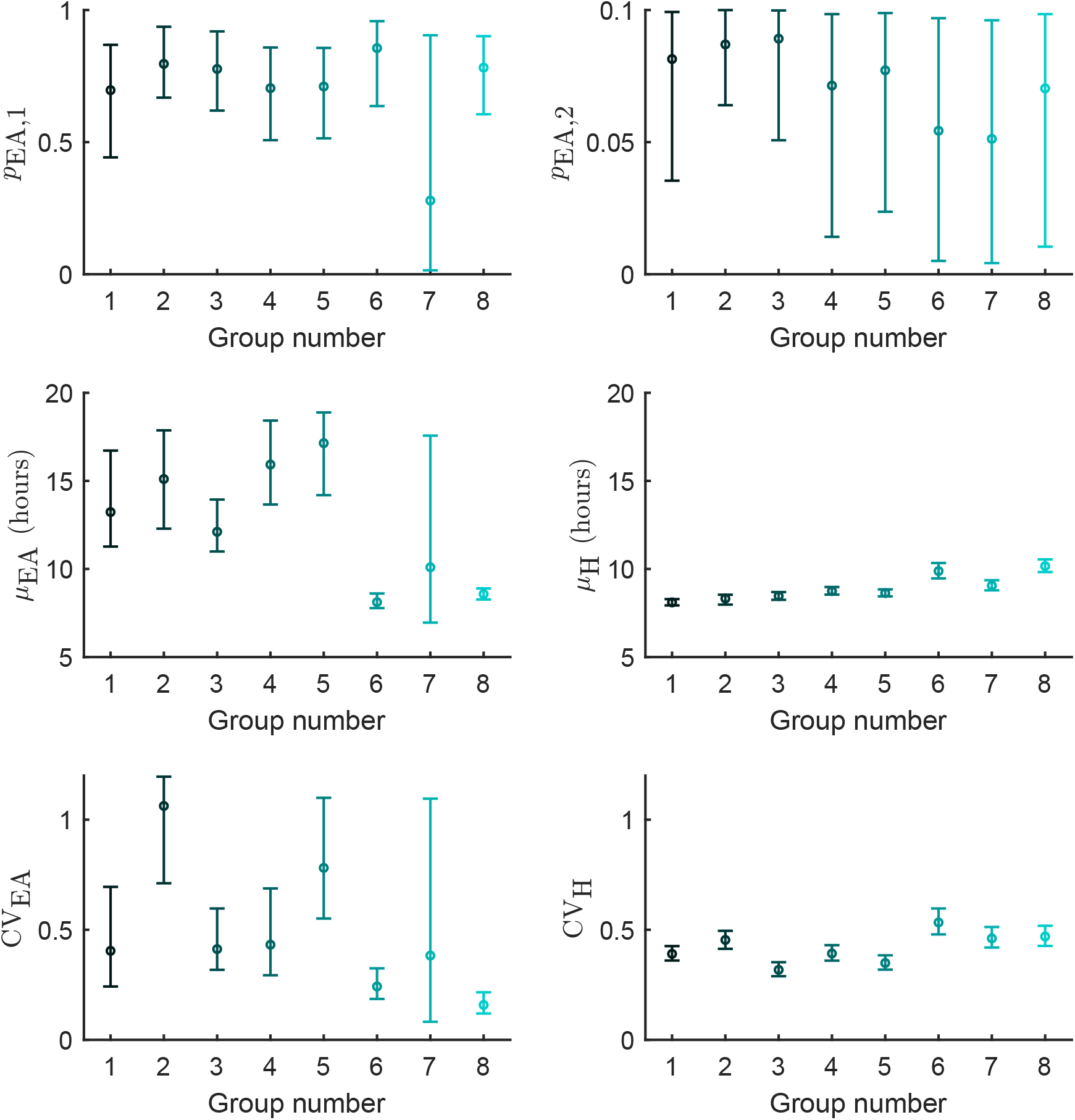
Inferred parameter values for model #2 in the eight data groups. The circles and the error bars respectively show the median and the 95% credible intervals of the parameter posterior distributions.

**Figure S19:**
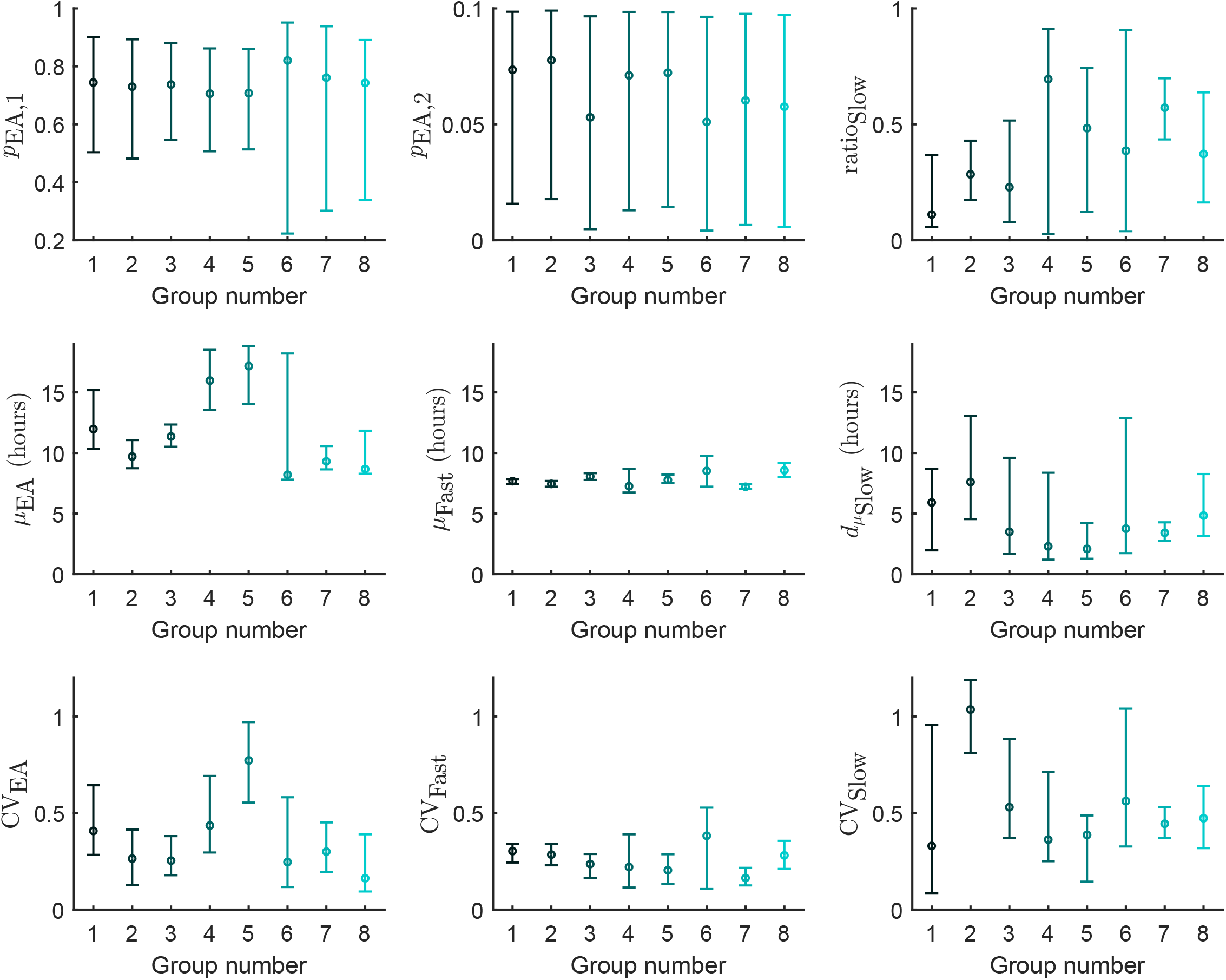
Inferred parameter values for the mixture model in the eight data groups. The circles and the error bars respectively show the median and the 95% credible intervals of the parameter posterior distributions.

**Figure S20:**
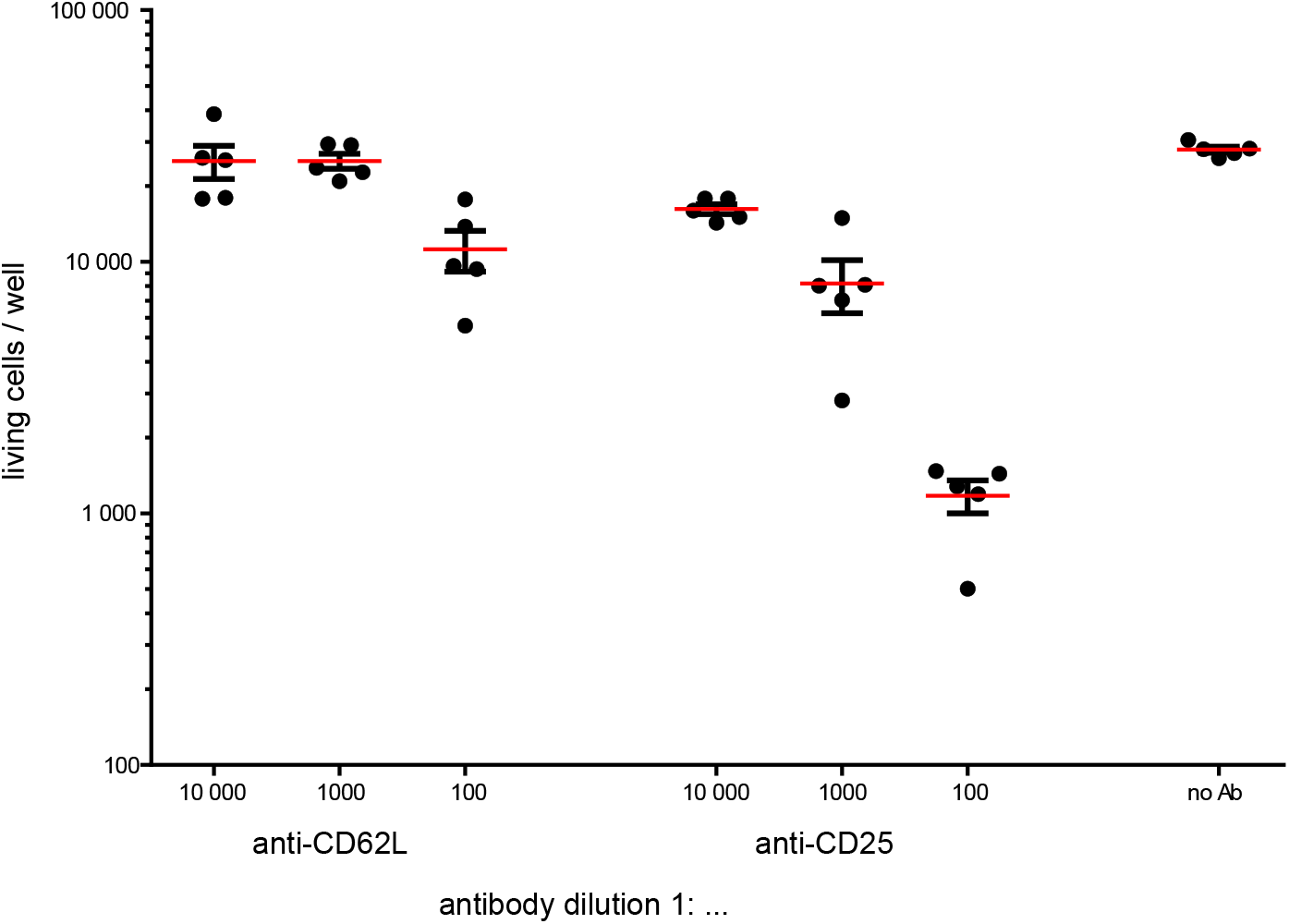
Impact of antibodies on proliferation *in vitro*. 100 naïve T cells were sorted per well (anti-CD3/CD28 coated). Anti-CD62L or anti-CD25 antibodies were added to the culture medium 1:100, 1:1000 or 1:10 000 diluted. After 5 days the cell number per well was measured using flow cytometry. Of note: For live cell imaging dilutions of 1:10 000 and 1:20 000 were used for anti-CD62L and anti-CD25, respectively.

**Figure S21:**
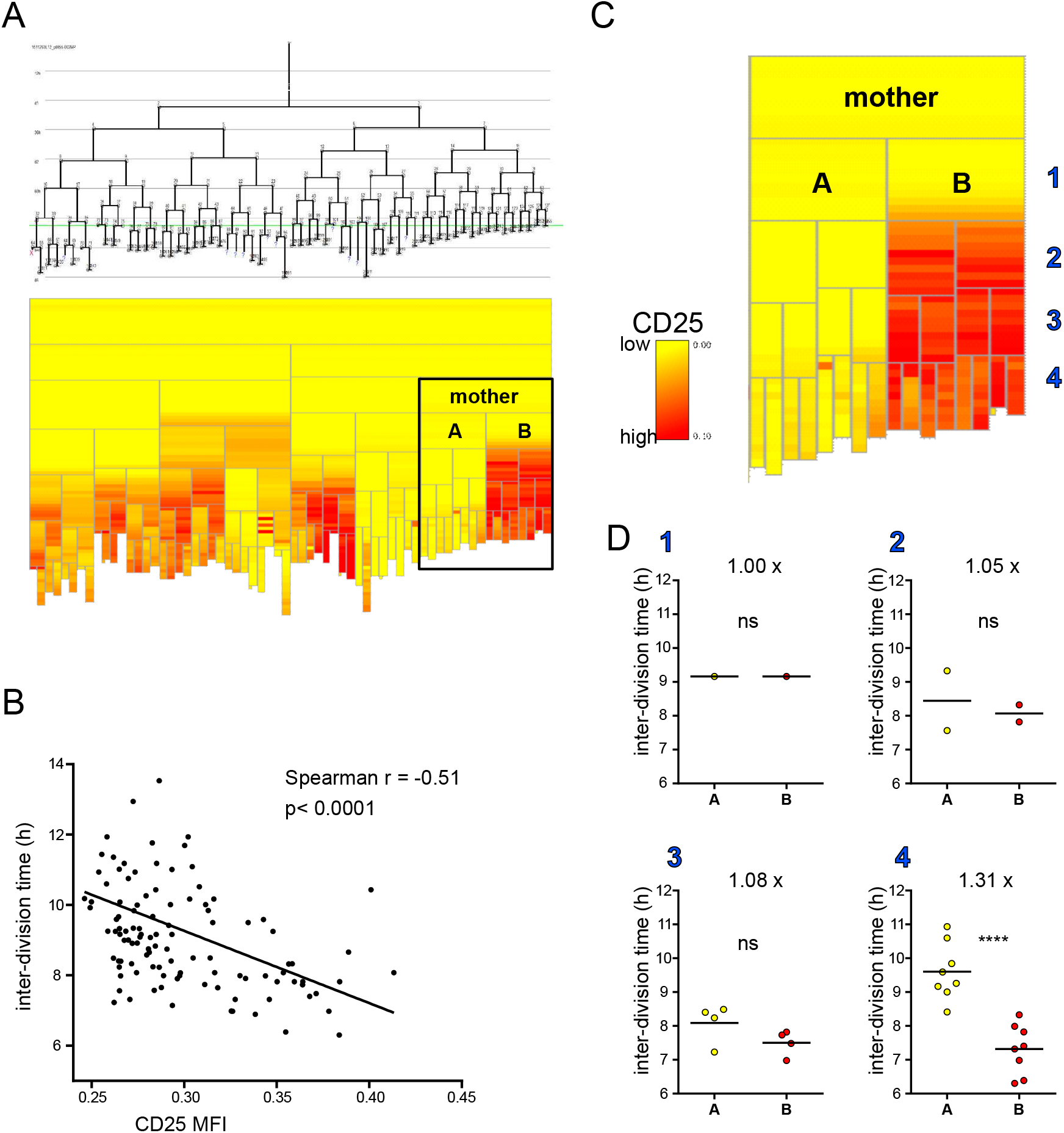
CD25 is enriched in distinct branches and correlates with division speed. Cells were stimulated with anti-CD3/CD28 and imaged in the presence of anti-CD25-APC. **(A)** A representative tree and its heattree plot as in Fig. 3F. Two very distinct (in terms of their CD25 expression) branches (A and B) and the mother cell from which these branches were derived are marked. **(B)** The inter-division times of all cells in (A) are plotted against their CD25 expression levels. **(C)** The cells in the marked branches from (A). Blue numbers (1–4) indicate the number of cell divisions starting from the division of the common mother cell. The inter-division times of these cells are compared within their respective generation (1-4, blue) between branch A and branch B **(D).** Numbers with x indicate the factor between the mean inter-division times of the cells in branches A and B. Student’s t-test: ****: p<0.0001.

**Figure S22:**
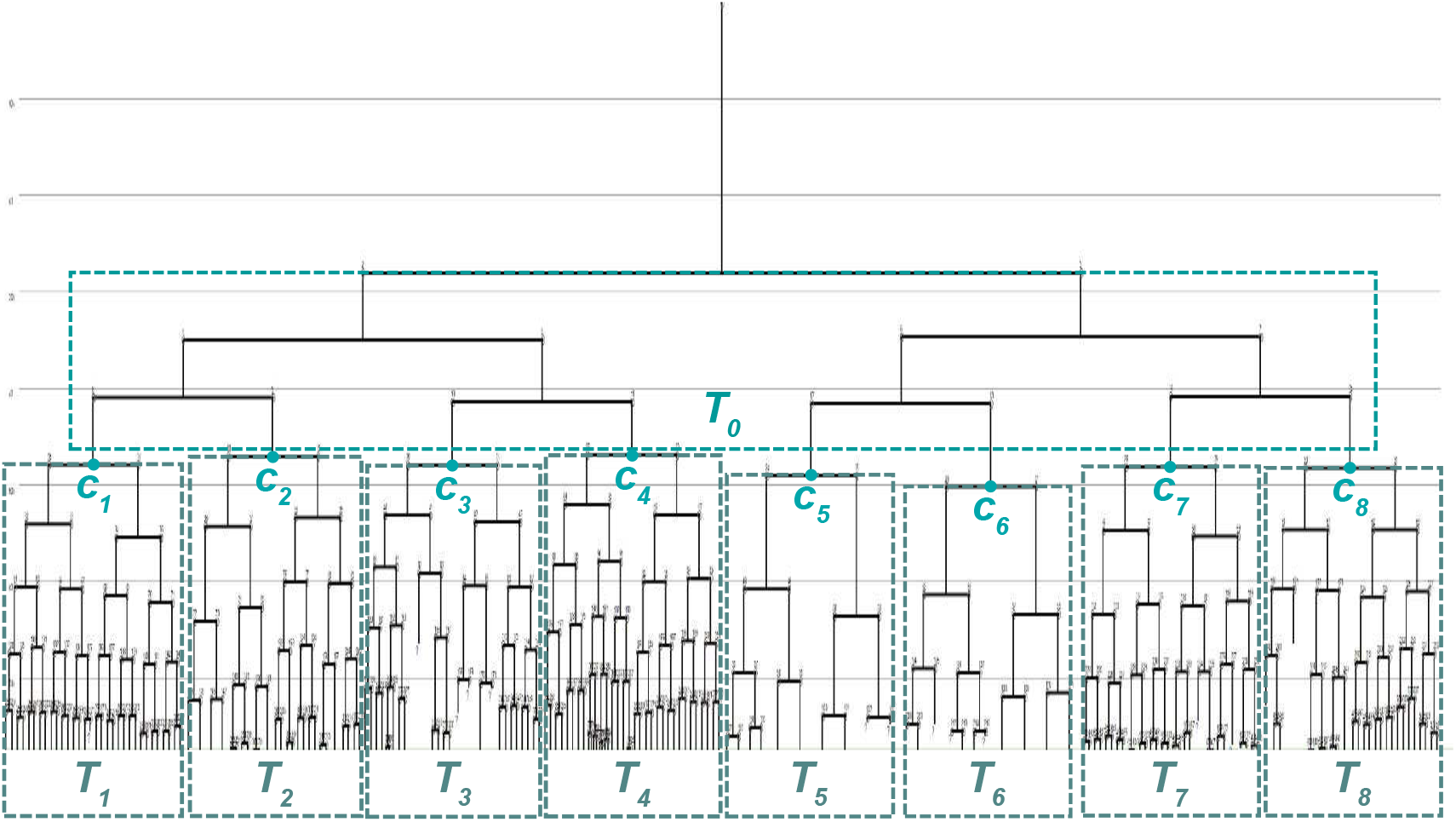
Modular division of a family tree for efficient sampling of latent variables. A family tree is divided into nine “subtrees”. The first subtree (*T*_0_) includes all cells belonging to generations 1, 2 and 3. The remaining subtrees (*T*_1_, *T*_2_,…, *T*_8_) are subtrees descending from the eight third-generation cells: *T*_1_ includes all cells descending from cell *c*_1_, *T*_2_ includes all cells descending from cell *c*_2_ and so forth.

**Movie S1:**

A single naïve OT-I cell was sorted on anti-CD3/CD28 (as in Fig. 1) and imaged for approximately four days. The numbers in the upper right corner indicate the time after start of the experiment (d – hh:mm:ss). The white circles or dots on the cells are virtual markers that were set manually to track the cells.

**Movie S2:**

A single activated OT-I cell was sorted into a “macrowell” without anti-CD3 coating and was imaged for approximately four days. Under these conditions, the T cells distributed over the complete well.

**Movie S3:**

A single naïve OT-I cell was sorted on anti-CD3/CD28 (as in Fig. 1) and imaged for approximately four days. The numbers in the upper right corner indicate the time after start of the experiment (d - hh:mm:ss). Every cell is marked with a colored tracing line to visualize the movement of the cells.

**Movie S4:**

A single naïve OT-I cell was sorted on anti-CD3/CD28 (as in Fig. 1) and imaged for approximately five days in the presence of anti-CD25-APC. The numbers in the upper right corner indicate the time after start of the experiment (d – hh:mm:ss). The bright dots on the cells are virtual markers that were set manually to track the cells. The bright field image is shown on the left side. The corresponding APC signal is shown on the right side. To reduce bleaching and phototoxicity, the APC channel was not acquired as often as the bright field. Thus, the APC channel does not always update together with the bright field image.

## Notes

### Competing Interest Statement

The authors have declared no competing interest.

## References

1. Burnet SFM (1959) The Clonal Selection Theory of Acquired Immunity (Cambridge University Press).

2. Buchholz VR, et al. (2013) Disparate individual fates compose robust CD8+ T cell immunity. Science 340(6132):630–635.

3. Gerlach C, et al. (2013) Heterogeneous differentiation patterns of individual CD8+ T cells. Science 340(6132):635–639.

4. Tubo NJ, et al. (2013) Single naive CD4+ T cells from a diverse repertoire produce different effector cell types during infection. Cell 153(4):785–796.

5. Plumlee CR, Sheridan BS, Cicek BB, Lefrançois L (2013) Environmental cues dictate the fate of individual CD8+ T cells responding to infection. Immunity 39(2):347–356.

6. Cho Y-L, et al. (2017) TCR Signal Quality Modulates Fate Decisions of Single CD4(+) T Cells in a Probabilistic Manner. Cell Rep 20(4):806–818.

7. Graef P, et al. (2014) Serial transfer of single-cell-derived immunocompetence reveals stemness of CD8(+) central memory T cells. Immunity 41(1):116–126.

8. Kretschmer L, et al. (2020) Differential expansion of T central memory precursor and effector subsets is regulated by division speed. Nat Commun 11(1): 113.

9. Lin W-HW, et al. (2016) CD8(+) T Lymphocyte Self-Renewal during Effector Cell Determination. Cell Rep 17(7):1773–1782.

10. Grassmann S, et al. (2020) Early emergence of T central memory precursors programs clonal dominance during chronic viral infection. Nat Immunol 170:2022.

11. Pais Ferreira D, et al. (2020) Central memory CD8+ T cells derive from stem-like Tcf7hi effector cells in the absence of cytotoxic differentiation. Immunity. doi:10.1016/j.immuni.2020.09.005.

12. Johnnidis JB, et al. (2021) Inhibitory signaling sustains a distinct early memory CD8+ T cell precursor that is resistant to DNA damage. Science Immunology 6(55):eabe3702.

13. Yoon H, Kim TS, Braciale TJ (2010) The cell cycle time of CD8+ T cells responding in vivo is controlled by the type of antigenic stimulus. PLoS ONE 5(11):e15423.

14. Kinjyo I, et al. (2015) Real-time tracking of cell cycle progression during CD8(+) effector and memory T-cell differentiation. Nat Commun 6:6301.

15. Hawkins ED, Markham JF, McGuinness LP, Hodgkin PD (2009) A single-cell pedigree analysis of alternative stochastic lymphocyte fates. Proceedings of the National Academy of Sciences 106(32):13457–13462.

16. Zaretsky I, et al. (2012) Monitoring the dynamics of primary T cell activation and differentiation using long term live cell imaging in microwell arrays. Lab Chip 12(23):5007–5015.

17. Polonsky M, Chain B, Friedman N (2016) Clonal expansion under the microscope: studying lymphocyte activation and differentiation using live-cell imaging. Immunol Cell Biol 94(3):242–249

18. Marchingo JM, et al. (2016) T-cell stimuli independently sum to regulate an inherited clonal division fate. Nat Commun 7:13540.

19. Hilsenbeck O, et al. (2016) Software tools for single-cell tracking and quantification of cellular and molecular properties. Nat Biotechnol 34(7):703–706.

20. Loeffler D, et al. (2019) Asymmetric lysosome inheritance predicts activation of haematopoietic stem cells. Nature 573(7774):426–429.

21. Harris T (1963) The Theory of Branching Processes. Springer, Berlin.

22. Wilkinson DJ (2009) Stochastic modelling for quantitative description of heterogeneous biological systems. Nat. Rev. Genet. 10:122–133.

23. Wilkinson DJ (2011) Stochastic Modelling for Systems Biology. Second Edition. CRC Press.

24. Eilken HM, Nishikawa S-I, Schroeder T (2009) Continuous single-cell imaging of blood generation from haemogenic endothelium. Nature 457(7231):896–900.

25. Moreau HD, et al. (2012) Dynamic in situ cytometry uncovers T cell receptor signaling during immunological synapses and kinapses in vivo. Immunity 37(2):351–363.

26. Stemberger C, et al. (2007) A single naive CD8+ T cell precursor can develop into diverse effector and memory subsets. Immunity 27(6):985–997.

27. Marchingo JM, et al. (2014) T cell signaling. Antigen affinity, costimulation, and cytokine inputs sum linearly to amplify T cell expansion. Science 346(6213):1123–1127.

28. Heinzel S, et al. (2017) A Myc-dependent division timer complements a cell-death timer to regulate T cell and B cell responses. Nat Immunol 18(1):96–103.

29. Arsenio J, et al. (2014) Early specification of CD8+ T lymphocyte fates during adaptive immunity revealed by single-cell gene-expression analyses. Nature Immunology 15(4):365–372.

30. Kakaradov B, et al. (2017) Early transcriptional and epigenetic regulation of CD8+ T cell differentiation revealed by single-cell RNA sequencing. Nature Immunology 18(4):422–432.

31. Chang JT, et al. (2007) Asymmetric T lymphocyte division in the initiation of adaptive immune responses. Science 315(5819):1687–1691.

32. Bird JJ, et al. (1998) Helper T cell differentiation is controlled by the cell cycle. Immunity 9(2):229–237.

33. Gett AV, Hodgkin PD (1998) Cell division regulates the T cell cytokine repertoire, revealing a mechanism underlying immune class regulation. Proc Natl Acad Sci USA 95(16):9488–9493.

34. Duffy K, et al. (2012) Activation-induced B cell fates are selected by intracellular stochastic competition. Science 279:338–341.

35. Marr C, et al. (2012) Multi-scale modeling of GMP differentiation based on single-cell genealogies. FEBS. J. 279:3488–3500.

36. Niederberger T, et al. (2015) Factor graph analysis of live cell imaging data reveals mechanisms of cell fate decisions. Bioinformatics 31:1816–1823.

37. Hormoz S, et al. (2016) Inferring cell-state transition dynamics from lineage trees and endpoint single-cell measurements. Cell Syst. 3:419–433.e8.

38. Feigelman J, et al. (2016) Analysis of Cell Lineage Trees by Exact Bayesian Inference Identifies Negative Autoregulation of Nanog in Mouse Embryonic Stem Cells. Cell Syst. 3(5):480–490.e13.

39. Strasser MK, et al. (2018) Lineage marker synchrony in hematopoietic genealogies refutes the PU.1/GATA1 toggle switch paradigm. Nat Commun 9:2697.

