## Supplementary Methods for "Heritable changes in division speed accompany the diversification of single T cell fate"

### 0. Introduction

We developed a Bayesian inference framework for hypothesis-driven analysis of cell lineage trees. This inference scheme tests different hypotheses to infer underlying pathways of diversification in single-cell-derived populations. For a given model hypothesis, it uses lineage (family) trees obtained from live-cell imaging experiments to infer the model parameters. We used this framework to test various hypotheses about the existence of subsets with distinct inter-division time statistics in T cell family trees expanded *in vitro* and the corresponding diversification pathways. The scheme presented here is specifically designed for bright-field microscopy imaging data where no direct phenotypic measurements are available. However, it can be generalized to analyze live-cell imaging with phenotypic marker expression data. In the following, we describe different components of our computational analysis. In Sections 1 and 2, we outline our Bayesian inference scheme and Bayesian model comparison. In Section 3, we discuss the details of different model hypotheses we tested for analyzing T cell family trees. Section 4 briefly outlines the statistical features of the data that we aim to capture with mathematical modeling. Finally, Section 5 describes the simulation studies that were used to examine how well mathematical models recapitulated the statistical properties of the data.

### 1. Bayesian model inference scheme for the analysis of live-cell imaging data

Our Bayesian framework utilizes a Markov Chain Monte Carlo (MCMC) scheme with latent variables (Wilkinson, 2009; Wilkinson, 2011) tailored to tree-structured data, incorporating familial relationships of individual cells within a family tree. This framework takes family trees obtained via live-cell imaging, as well as a model hypothesis, as input and returns the posterior distribution of model parameters as output. It enables model comparison and hypothesis testing through the approximation of model evidences and corresponding Bayes factors. Different steps of our inference scheme are described in the following.

**1.1. Input data.** Family trees generated by The Tracking Tool (Hilsenbeck et al., 2016) from live-cell imaging experiments form the input data of the inference scheme (see Materials and Methods of the main text). This data encodes two types of information for every cell in a family tree: 1) the lifetime of that cell, and 2) the index of the cell within the family tree which yields familial relationships. Dataset $D$ consists of the lifetimes of individual cells across different trees:

$$D = \{\tilde{t}_{im} | i = 1, \dots, n_{\text{tree}}, m = 1, \dots, n_{\text{cell}_i}\} \tag{1.1}$$

where $n_{\text{tree}}$ is the total number of trees and $n_{\text{cell}_i}$ is the total number of cells in tree $i$ and $\tilde{t}_{im}$ is the lifetime of cell $m$ in tree $i$. If cell $m$ has divided in the course of the experiment, $\tilde{t}_{im}$ denotes its inter-division time; otherwise, it represents the time until which this cell had not divided indicating $\tilde{t}_{im}$ as a lower bound for the inter-division time. We assume that the interdivision times are log-normally distributed independent random variables. We further assume a minimum cell-cycle length $\tau$ to exclude un-physiologically short inter-division times. To enable robust numerical calculations, we first subtract the threshold $\tau$ and then log-transform the data in $D$ to arrive at

$$\boldsymbol{y} = \left\{t_{im} \middle| i = 1, \dots, n_{\text{tree}}, m = 1, \dots, n_{\text{cell}_i}\right\}, \quad \text{with } t_{im} = \log\left(\tilde{t}_{im} - \tau\right) \tag{1.2}$$

The dataset $\boldsymbol{y}$ is then used to infer the model structure and parameters.

**1.2. Input model hypothesis.** The inference scheme relies on a model hypothesis for the computational analysis of the data. This model hypothesis represents assumptions about the existence of distinct subsets in the population and the diversification pathways connecting these subsets. We used our inference scheme to analyze family trees in the absence of surface marker expression measurements. Therefore, no subsets defined based on marker combinations were considered in this study. Instead, subsets of cells which possess distinct inter-division time statistics were considered. However, if direct phenotypic measurements are available, they can be readily incorporated into the model hypothesis and the inference scheme.

In Section 3, we provide a detailed description of model topologies considered in this study. In all of the hypotheses, we assume that the inter-division times of cells belonging to a specific subset follow a log-normal distribution with specific but unknown mean and coefficient of variation. Other distribution assumptions can easily replace the lognormal assumption, and can be handled in the inference scheme the same way. Each model hypothesis is characterized by a set of unknown model parameters $\boldsymbol{\theta}$ such as the mean and coefficient of variation of the inter-division time distributions as well as the transition probabilities between different subsets. Furthermore, the subset that is assigned to a cell is modeled as a latent variable due to the unavailability of direct observation.

**1.3. Bayesian Inference scheme.** We use a Markov Chain Monte Carlo (MCMC) approach to generate samples from the posterior distribution of the model parameters $\boldsymbol{\theta}$ given the data $\boldsymbol{y}$. For all parameters, we assume uniform prior distributions with specified upper and lower bounds. Different steps of the iterative sampling scheme are described in the following sections, and a pseudo-code is given in Algorithm 1.

**1.3.1. MCMC sampling for models with latent variables.** As mentioned earlier, the underlying subset that a cell in the family tree belongs to, is not directly observed and therefore needs to be modeled as a latent variable. For a model with latent variables, we have the following factorization

$$\pi(\boldsymbol{\theta}, \boldsymbol{x}, \boldsymbol{y}) = \pi(\boldsymbol{\theta})\pi(\boldsymbol{x}|\boldsymbol{\theta})\pi(\boldsymbol{y}|\boldsymbol{x}, \boldsymbol{\theta}) \tag{1.3}$$

where, $\boldsymbol{\theta}$ is the vector of model parameters, $\boldsymbol{x}$ is the vector of latent variables and $\boldsymbol{y}$ is the data. Here, $\boldsymbol{x}$ and $\boldsymbol{y}$ are respectively the assigned subsets and the lifetimes of individual cells. Since we are only interested in the posterior distribution of the model parameters $\pi(\boldsymbol{\theta}|\boldsymbol{y})$, we can use an arbitrary proposal $q(\boldsymbol{\theta}, \boldsymbol{\theta}^*)$, to target this posterior using the following acceptance ratio:

$$A = \frac{\pi(\boldsymbol{\theta}^*)\ \pi(\boldsymbol{y}|\boldsymbol{\theta}^*)\ q(\boldsymbol{\theta}^*, \boldsymbol{\theta})}{\pi(\boldsymbol{\theta})\ \pi(\boldsymbol{y}|\boldsymbol{\theta})\ q(\boldsymbol{\theta}, \boldsymbol{\theta}^*)} \tag{1.4}$$

In order to calculate the marginal likelihood of the data given the parameters $\pi(\boldsymbol{y}|\boldsymbol{\theta})$, we must integrate out the latent variables:

$$\pi(\boldsymbol{y}|\boldsymbol{\theta}) = \int_{\boldsymbol{X}} \pi(\boldsymbol{y}|\boldsymbol{x}, \boldsymbol{\theta})\ \pi(\boldsymbol{x}|\boldsymbol{\theta})\ d\boldsymbol{x} \tag{1.5}$$

Due to the intractability of the analytical calculation of this integral, we aim to use a Monte Carlo approximation of it

$$\hat{\pi}(\boldsymbol{y}|\boldsymbol{\theta}) = \frac{1}{n_x} \sum_{k=1}^{n_x} \pi(\boldsymbol{y}|\boldsymbol{x}_k, \boldsymbol{\theta}) \tag{1.6}$$

It can be shown that since this is an unbiased estimator of the likelihood, if we plug this approximation into the acceptance ratio (Eq. 1.4), we will exactly target the posterior distribution $\pi(\boldsymbol{\theta}|\boldsymbol{y})$ (Wilkinson, 2011):

$$A = \frac{\pi(\boldsymbol{\theta}^*)\ \ \hat{\pi}(\boldsymbol{y}|\boldsymbol{\theta}^*)\ q(\boldsymbol{\theta}^*, \boldsymbol{\theta})}{\pi(\boldsymbol{\theta})\ \ \hat{\pi}(\boldsymbol{y}|\boldsymbol{\theta})\ q(\boldsymbol{\theta}, \boldsymbol{\theta}^*)} \tag{1.7}$$

Here, $\hat{\pi}(\boldsymbol{y}|\boldsymbol{\theta})$ denotes our unbiased estimate of the marginal likelihood $\pi(\boldsymbol{y}|\boldsymbol{\theta})$ obtained by the Monte Carlo integration above. We assume symmetric prior and proposal distributions, which cancels out the terms $\pi(\boldsymbol{\theta}^*)\ q(\boldsymbol{\theta}^*, \boldsymbol{\theta})$ and $\pi(\boldsymbol{\theta})\ q(\boldsymbol{\theta}, \boldsymbol{\theta}^*)$ in the numerator and denominator of Eq. (1.7), resulting in the acceptance ratio

$$A = \frac{\hat{\pi}(\boldsymbol{y}|\boldsymbol{\theta}^*)}{\hat{\pi}(\boldsymbol{y}|\boldsymbol{\theta})} \tag{1.8}$$

We propose parameter sets according to the proposal $q(\boldsymbol{\theta}, \boldsymbol{\theta}^*)$, and based on the acceptance ratio Eq. 1.8, obtain samples from the posterior distribution $\pi(\boldsymbol{\theta}|\boldsymbol{y})$. In the study of T cell family trees, we used random walk and Gaussian proposals.

**1.3.2. Unbiased estimates of the likelihood $\hat{\pi}(\boldsymbol{y}|\boldsymbol{\theta}^*)$.** We use the following Monte Carlo approximation based on the latent variable samples to obtain an unbiased estimate of the likelihood:

$$\hat{\pi}(\boldsymbol{y}|\boldsymbol{\theta}) = \frac{1}{n_x} \sum_{k=1}^{n_x} \pi(\boldsymbol{y}|\boldsymbol{x}_k, \boldsymbol{\theta}) \tag{1.9}$$

Since our data is comprised of several lineage trees, which are independent from each other, the data likelihood $\pi(\boldsymbol{y}|\boldsymbol{\theta})$ can be written as

$$\pi(\boldsymbol{y}|\boldsymbol{\theta}) = \prod_{i=1}^{n_{\text{tree}}} \pi(\boldsymbol{y}_i|\boldsymbol{\theta}) \tag{1.10}$$

where $\boldsymbol{y}_i$ is the data belonging to the $i^{\text{th}}$ lineage tree and $\pi(\boldsymbol{y}_i|\boldsymbol{\theta})$ is its likelihood. Moreover, the latent variables of every tree, i.e. the subsets of cells within that tree, are independent from other trees. Thus, we can apply the Monte Carlo approximation on every tree individually and reconstruct an unbiased estimate of the overall likelihood based on individual tree likelihoods. In this way we have

$$\hat{\pi}(\boldsymbol{y}|\boldsymbol{\theta}) = \prod_{i=1}^{n_{\text{tree}}} \hat{\pi}(\boldsymbol{y}_i|\boldsymbol{\theta}) \tag{1.11}$$

where $\hat{\pi}(\boldsymbol{y}_i|\boldsymbol{\theta})$ is the unbiased estimate of the likelihood of the $i^{\text{th}}$ tree. For numerical properties, it is favorable to work with the log-likelihood. Taking the log-transformation of Eq. 1.11, we obtain the following for the log-likelihood:

$$\hat{J}(\boldsymbol{\theta}) = \log\left(\hat{\pi}(\boldsymbol{y}|\boldsymbol{\theta})\right) = \sum_{i=1}^{n_{\text{tree}}} \log(\hat{\pi}(\boldsymbol{y}_i|\boldsymbol{\theta})). \tag{1.12}$$

We next need to sample latent variables in every tree to calculate the log-likelihoods $\log(\hat{\pi}(\boldsymbol{y}_i|\boldsymbol{\theta}))$.

**1.3.3. Two-step sampling of the latent variables.** To construct an unbiased estimate of the likelihood of a tree given the parameters, $\hat{\pi}(\boldsymbol{y}_i|\boldsymbol{\theta})$, we generate several samples of the latent variables, for each of which the likelihood can be directly evaluated. The number of possible configurations for latent variables increases exponentially with the number of cells. To increase the efficiency and the acceptance ratio of our sampling scheme, we divide each tree into nine "subtrees" (Fig. S22). We perform sample generation and likelihood approximation for these subtrees in a modular fashion. The first subtree, denoted as $T_0$, includes all cells belonging to generations 1, 2 and 3. The remaining subtrees, denoted as $T_1, T_2, \ldots, T_8$, are subtrees descending from the eight third-generation cells: $T_1$ includes all cells descending from cell $c_1$, $T_2$ includes all cells descending from cell $c_2$ and so forth (Fig. S22). In this way, cells $c_1, c_2, \ldots, c_8$, which themselves belong to $T_0$, form the founder cells for subtrees $T_1, T_2, \ldots, T_8$. The dependency between these subtrees is taken into account in the generation of the latent variable samples. Therefore, this modular division of the tree does not result in any loss of information and ensures that the likelihood of the whole tree is calculated at once. Subtree $T_0$—comprising of three cell divisions—is generally smaller than subtrees $T_1, T_2, \ldots, T_8$ which can comprise of arbitrarily many cell divisions (in our data up to six divisions). Thus, there are considerably fewer possible configurations for $T_0$ than for $T_1, T_2, \ldots, T_8$. To exploit this property for more efficient sampling, we propose a two-step sampling strategy: 1) first a set of latent samples for $T_0$ is generated and 2) based on each single sample, a second set of latent samples for $T_1, T_2, \ldots, T_8$ is generated. In this way, the configuration of subtrees $T_1, T_2, \ldots, T_8$ is sampled more often than $T_0$, increasing the chance of generating high-likelihood samples and thereby increasing the acceptance ratio of the sampling approach. This division of the tree is further encouraged by our data: in the T cell family trees, we observed an initial burst phase of semi-concordant cell divisions (corresponding to $T_0$), which is followed by more variable division speeds in the subsequent divisions (corresponding to $T_1, T_2, \ldots, T_8$) (see the main text). This correspondence facilitates using different parameters for describing the first division phase ($T_0$) and the subsequent division phase ($T_1, T_2, \ldots, T_8$), and lets us tune the adequate number of latent samples in each phase separately.

In the first step, we generate $ns_1$ samples of latent variables for all cells in subtree $T_0$. We denote the set of latent variables for $T_0$ by $\boldsymbol{x}_k^0$:

$$\boldsymbol{x}_k^0 = \left\{ x_k^{0,m} \middle| m = 1, \ldots, n_{T_0} \right\} \quad \text{for} \quad k = 1, \ldots, ns_1 \tag{1.13}$$

where $n_{T_0}$ is the number of cells in subtree $T_0$. As explained in Section 3, we use a branching process framework (Harris, 1963) to randomly simulate latent variables $x_k^{0,m}$ according to the assumed model topology and its parameters. In this way, the latent variable of every cell is only dependent on the latent variable of its mother cell. Therefore, the configuration of subtree $T_0$, $\boldsymbol{x}_k^0$, is independent from the configuration of subtrees $T_1, T_2, \ldots, T_8$. On the contrary, the latent states of $T_1, T_2, \ldots, T_8$ depend on the their corresponding founder cells $c_1, c_2, \ldots, c_8$.

In the second step, for each latent sample of $c_1, c_2, \ldots, c_8$, we generate $ns_2$ latent samples for subtrees $T_1, T_2, \ldots, T_8$. We denote the latent states of subtree $T_j$ by $\boldsymbol{x}_{k,l}^j$:

$$\boldsymbol{x}_{k,l}^j = \left\{ x_{k,l}^{j,m} \middle| m = 1, \ldots, n_{T_j} \right\} \quad \text{for} \quad k = 1, \ldots, ns_1, \quad l = 1, \ldots, ns_2, \quad j = 1, \ldots, 8 \tag{1.14}$$

where $n_{T_j}$ is the number of cells in subtree $T_j$. Each latent sample $\boldsymbol{x}_{k,l}^j$ for subtree $T_j$ depends on the latent sample $\boldsymbol{x}_k^0$ of subtree $T_0$ through the state of the founder cells $c_1, c_2, \ldots, c_8$. Since the lifetimes of individual cells are assumed to be independent random variables, the only dependence between subtrees is through topological dependence of latent variables. Thus, once latent variables are sampled, the data of different subtrees are independent from each other. For a given latent variable sample $\boldsymbol{x}^*$, the likelihood of the whole tree can be written as

$$\pi(\boldsymbol{y}|\boldsymbol{x}^*, \boldsymbol{\theta}) = \prod_{j=0}^{8} \pi\left(\boldsymbol{y}^j \; |\boldsymbol{x}^*, \boldsymbol{\theta}\right) \tag{1.15}$$

For the simplicity of notation, we have dropped the index of tree in this section and use $\boldsymbol{y}$ to denote the data belonging to a given tree. $\boldsymbol{y}^j$ represents the data corresponding to subtree $T_j$. We then use the following Monte Carlo approximation to integrate out the two-layer latent variable samples and estimate the tree likelihood given the parameters:

$$\hat{\pi}(\boldsymbol{y}|\boldsymbol{\theta}) = \frac{1}{ns_1} \sum_{k=1}^{ns_1} \hat{\pi}(\boldsymbol{y}|\boldsymbol{x}_k^0, \boldsymbol{\theta}) = \frac{1}{ns_1} \sum_{k=1}^{ns_1} \left( \pi\left(\boldsymbol{y}^0 | \boldsymbol{x}_k^0, \boldsymbol{\theta}\right) \prod_{j=1}^{8} \hat{\pi}\left(\boldsymbol{y}^j | \boldsymbol{x}_k^0, \boldsymbol{\theta}\right) \right)$$
$$= \frac{1}{ns_1} \sum_{k=1}^{ns_1} \left( \pi\left(\boldsymbol{y}^0 | \boldsymbol{x}_k^0, \boldsymbol{\theta}\right) \prod_{j=1}^{8} \frac{1}{ns_2} \sum_{l=1}^{ns_2} \pi\left(\boldsymbol{y}^j | \boldsymbol{x}_{k,l}^j, \boldsymbol{\theta}\right) \right) \tag{1.16}$$

As we need $\log(\hat{\pi}(\boldsymbol{y}|\boldsymbol{\theta}))$ for the calculation of overall data log-likelihood in Eq. 1.12, we take the logarithm of the equation above. For better readability, we drop the dependence on $\boldsymbol{\theta}$ in the rest of the equations in this section, however it is implied that all the data likelihoods are conditioned on the parameter set $\boldsymbol{\theta}$:

$$\log(\hat{\pi}(\boldsymbol{y})) = \log\left(\frac{1}{ns_1}\right) + \log\left( \sum_{k=1}^{ns_1} \left( \pi\left(\boldsymbol{y}^0 | \boldsymbol{x}_k^0\right) \prod_{j=1}^{8} \frac{1}{ns_2} \sum_{l=1}^{ns_2} \pi\left(\boldsymbol{y}^j | \boldsymbol{x}_{k,l}^j\right) \right) \right) \tag{1.17}$$

The logarithm cannot be pushed inside the sum in the second term of Eq. 1.17; therefore, we use the following numerical trick to robustly calculate this term based on the individual subtree log-likelihoods. More details about this robust numerical evaluation can be found in (Loos, 2016; Loos et al., 2016). We define $q_k$ as the logarithm of the summand in the second term of Eq. 1.17 :

$$q_k = \log\left( \pi\left(\boldsymbol{y}^0 | \boldsymbol{x}_k^0\right) \prod_{j=1}^{8} \frac{1}{ns_2} \sum_{l=1}^{ns_2} \pi\left(\boldsymbol{y}^j | \boldsymbol{x}_{k,l}^j\right) \right) \tag{1.18}$$

$q^*$ denotes the maximum of $q_k$:

$$q^* = \arg\max_k q_k \tag{1.19}$$

The second term in Eq. 1.17 can then be evaluated via

$$\log\left( \sum_{k=1}^{ns_1} \left( \pi\left(\boldsymbol{y}^0 | \boldsymbol{x}_k^0\right) \prod_{j=1}^{8} \frac{1}{ns_2} \sum_{l=1}^{ns_2} \pi\left(\boldsymbol{y}^j | \boldsymbol{x}_{k,l}^j\right) \right) \right) = \log\left( \sum_{k=1}^{ns_1} e^{(q_k - q^*)} \right) + \log(q^*) \tag{1.20}$$

Inserting 1.20 into 1.17, we obtain the following for the log-likelihood of a tree:

$$\log(\hat{\pi}(\boldsymbol{y})) = \log\left(\frac{1}{ns_1}\right) + \log\left( \sum_{k=1}^{ns_1} e^{(q_k - q^*)} \right) + \log(q^*) \tag{1.21}$$

According to Eq. 1.18, $q_k$ is given as

$$q_k = \log\left(\pi\left(\boldsymbol{y}^0 \middle| \boldsymbol{x}_k^0\right)\right) + 8 \log\left(\frac{1}{ns_2}\right) + \sum_{j=1}^{8} \log\left(\sum_{l=1}^{ns_2} \pi\left(\boldsymbol{y}^j \middle| \boldsymbol{x}_{k,l}^j\right)\right) \tag{1.22}$$

To calculate the summands of the last term in Eq. 1.22, we use the same numerical trick as above. We define $p_l^j$ as

$$p_l^j = \log\left(\pi\left(\boldsymbol{y}^j \middle| \boldsymbol{x}_{k,l}^j\right)\right) \tag{1.23}$$

and denote the maximum of $p_l^j$ as $p^{*,j}$:

$$p^{*,j} = \arg\max_l p_l^j \tag{1.24}$$

We then evaluate the logarithm of the summation in Eq. 1.22 as:

$$\log\left(\sum_{l=1}^{ns_2} \pi\left(\boldsymbol{y}^j \middle| \boldsymbol{x}_{k,l}^j\right)\right) = \log\left(\sum_{l=1}^{ns_2} e^{\left(p_l^j - p^{*,j}\right)}\right) + \log\left(p^{*,j}\right) \tag{1.25}$$

Inserting Eq. 1.25 into Eq. 1.22, we obtain

$$q_k = \log\left(\pi\left(\boldsymbol{y}^0 \middle| \boldsymbol{x}_k^0\right)\right) + 8 \log\left(\frac{1}{ns_2}\right) + \sum_{j=1}^{8} \left(\log\left(\sum_{l=1}^{ns_2} e^{\left(p_l^j - p^{*,j}\right)}\right) + \log\left(p^{*,j}\right)\right) \tag{1.26}$$

To evaluate the log-likelihood for a given tree based on Equations 1.21, 1.23 and 1.26, we finally need to calculate $\log\left(\pi\left(\boldsymbol{y}^0 \middle| \boldsymbol{x}_k^0\right)\right)$ and $\log\left(\pi\left(\boldsymbol{y}^j \middle| \boldsymbol{x}_{k,l}^j\right)\right)$. For every latent sample $\boldsymbol{x}_k^0$, we calculate the log-likelihood of subtree $T_0$ by:

$$J_k^0 = \log\left(\pi\left(\boldsymbol{y}^0 \middle| \boldsymbol{x}_k^0\right)\right) = \sum_{m=1}^{n_{T_0}} \log\left(\pi\left(t_m^0 \middle| x_k^{0,m}\right)\right) = \sum_{m=1}^{n_{T_0}} J_k^{0,m} \tag{1.27}$$

Similarly, for each latent sample $\boldsymbol{x}_{k,l}^j$ dependent on $\boldsymbol{x}_k^0$, we calculate the log-likelihood of subtree $T_j$ by:

$$J_{k,l}^j = \log\left(\pi\left(\boldsymbol{y}^j \middle| \boldsymbol{x}_{k,l}^j\right)\right) = \sum_{m=1}^{n_{T_j}} \log\left(\pi\left(t_m^j \middle| x_{k,l}^{j,m}\right)\right) = \sum_{m=1}^{n_{T_j}} J_{k,l}^{j,m} \tag{1.28}$$

The likelihood of individual cell lifetimes $\pi\left(t_m^0 \middle| x_k^{0,m}\right)$ and $\pi\left(t_m^j \middle| x_{k,l}^{j,m}\right)$ are determined based on the model hypothesis. The likelihoods for the models tested in this study are described in Section 3.

**1.4. Output.** This inference scheme returns samples from the posterior distribution of parameters $\pi(\boldsymbol{\theta}|\boldsymbol{y})$ as output. It also provides the corresponding likelihood value for every sampled parameter set. The latter is used for the calculation of model evidence and model comparison.

### 2. Bayesian model comparison.

To compare the ability of different model hypotheses for explaining the lineage tree data, we perform Bayesian model comparison based on Bayes factors. We use the *Posterior Harmonic Mean estimator* (Vyshemirsky et al., 2007) to approximate the likelihood of the data given a model hypothesis $M$:

$$\pi(\boldsymbol{y}|M) \approx \left(\frac{1}{N}\sum_{i=1}^{N}\frac{1}{\pi\left(\boldsymbol{y}\middle|M,\boldsymbol{\theta}^{(i)}\right)}\right)^{-1}, \qquad \boldsymbol{\theta}^{(i)} \sim \pi(\boldsymbol{\theta}|\boldsymbol{y},M) \tag{2.1}$$

where $\boldsymbol{\theta}^{(i)}$ are samples from the parameter posterior and $\pi\left(\boldsymbol{y}\middle|M,\boldsymbol{\theta}^{(i)}\right)$ are the corresponding likelihoods. We approximate the Bayes factor comparing model $M_1$ to model $M_2$ using the model evidences:

$$B_{12} = \frac{\pi(\boldsymbol{y}|M_1)}{\pi(\boldsymbol{y}|M_2)} \tag{2.2}$$

To quantify the uncertainty of these approximation, we drew 10,000 sets of samples from the parameter posterior distribution of every model. This yielded a distribution for the approximated model evidences and Bayes factors.

**Algorithm 1:** Bayesian inference scheme for tree-structured data

**Data:** A set of single cell lifetimes $\boldsymbol{y} = \{t_{im} | i = 1, \dots, n_{\text{tree}}, m = 1, \dots, n_{\text{cell}_i}\}$ from $n_{\text{tree}}$ lineage trees each having $n_{\text{cell}_i}$ cells; a model hypothesis; parameter prior $\pi(\boldsymbol{\theta})$; number of iterations $N$

**Results:** A set of samples $\{\boldsymbol{\theta}^{(n)}\}_{n=1}^{N}$ from the posterior distribution of parameters $\pi(\boldsymbol{\theta}|\boldsymbol{y})$

Initialization;

Sample a set of parameter values from the prior: $\boldsymbol{\theta}^{(0)} \sim \pi(\boldsymbol{\theta})$;

Calculate the log-likelihood of the data given the initial parameter set: $\hat{J}^{(0)} = \log\left(\hat{\pi}\left(\boldsymbol{y}\middle|\boldsymbol{\theta}^{(0)}\right)\right)$

Initialize the chain of sampled parameter values and the corresponding log-likelihood values: $\boldsymbol{\theta}_{\text{chain}} \coloneqq \left[\boldsymbol{\theta}^{(0)}\right],\ J_{\text{chain}} \coloneqq [\,\hat{J}^{(0)}\,]$;

Iterate over the parameter samples;

**for** $n = 1 \dots N$ **do**

Generate a proposed set of parameter values $\boldsymbol{\theta}^*$ using the transition kernel (proposal distribution) $q\left(\boldsymbol{\theta}^{(n-1)}, \boldsymbol{\theta}^*\right)$: $\boldsymbol{\theta}^* \sim q\left(\boldsymbol{\theta}^{(n-1)}, \boldsymbol{\theta}^*\right) = f(\boldsymbol{\theta}^*|\boldsymbol{\theta}^{(n-1)})$.

**for** $i = 1 \dots n_{\text{tree}}$ **do**

**for** $k = 1 \dots ns_1$ **do**

Simulate a sample of latent variables of all cells in subtree $T_0$, according to the model topology and proposed parameters:

$\boldsymbol{x}_k^0 = \{x_k^{0,m} | m = 1, \dots, n_{T_0}\}$

Calculate the log-likelihood of subtree $T_0$ given $\boldsymbol{x}_k^0$ and the proposed parameter values $\boldsymbol{\theta}^*$:

$$J_k^0 = \log\left(\pi\left(\boldsymbol{y}_i^0\middle|\boldsymbol{x}_k^0, \boldsymbol{\theta}^*\right)\right) = \sum_{m=1}^{n_{T_0}} \log\left(\pi\left(t_m^0\middle|x_k^{0,m}, \boldsymbol{\theta}^*\right)\right) = \sum_{m=1}^{n_{T_0}} J_k^{0,m}$$

**for** $l = 1 \dots ns_2$ **do**

**for** $j = 1 \dots 8$ **do**

Simulate a sample of latent variables of all cells in subtree $T_j$, according to the model topology and proposed parameters:

$\boldsymbol{x}_{k,l}^j = \left\{x_{k,l}^{j,m} \middle| m = 1, \dots, n_{T_j}\right\}$

Calculate the log-likelihood of subtree $T_j$ given $\boldsymbol{x}_{k,l}^j$ and the proposed parameter values $\boldsymbol{\theta}^*$:

$$p_l^j = J_{k,l}^j = \log\left(\pi\left(\boldsymbol{y}_i^j \middle| \boldsymbol{x}_{k,l}^j, \boldsymbol{\theta}^*\right)\right)$$

$$= \sum_{m=1}^{n_{T_j}} \log\left(\pi\left(t_m^j \middle| x_{k,l}^{j,m}, \boldsymbol{\theta}^*\right)\right) = \sum_{m=1}^{n_{T_j}} J_{k,l}^{j,m}$$

Calculate the log-likelihood of tree $i$ given the latent variable given $\boldsymbol{x}_k^0$ and the proposed parameter values $\boldsymbol{\theta}^*$:

$$q_k = J_k^0 + 8\log\left(\frac{1}{ns_2}\right) + \sum_{j=1}^{8}\left(\log\left(\sum_{l=1}^{ns_2} e^{(p_l^j - p^{*,j})}\right) + \log\left(p^{*,j}\right)\right)$$

Calculate the log-likelihood of tree $i$ given the proposed parameter values $\boldsymbol{\theta}^*$:

$$\hat{J}_i = \log\left(\hat{\pi}(\boldsymbol{y}_i|\boldsymbol{\theta}^*)\right) = \log\left(\frac{1}{ns_1}\right) + \log\left(\sum_{k=1}^{ns_1} e^{(q_k - q^*)}\right) + \log(q^*)$$

Calculate the overall data log-likelihood given the proposed parameter values $\boldsymbol{\theta}^*$:

$$\hat{J}^* = \hat{\pi}(\boldsymbol{y}|\boldsymbol{\theta}^*) = \exp\left(\sum_{i=1}^{n_{\text{tree}}} \hat{J}_i\right)$$

Evaluate the acceptance probability of the proposed parameter values $\boldsymbol{\theta}^*$**:**

$$\alpha\left(\boldsymbol{\theta}^{(n-1)}, \boldsymbol{\theta}^*\right) = \min\left\{1, \frac{\hat{\pi}(\boldsymbol{y}|\boldsymbol{\theta}^*)}{\hat{\pi}\left(\boldsymbol{y}\middle|\boldsymbol{\theta}^{(n-1)}\right)}\right\} = \min\left\{1, \frac{\hat{J}^*}{\hat{J}^{(n-1)}}\right\}.$$

**If** uniform random variable $r \sim U(0,1) < \alpha\left(\boldsymbol{\theta}^{(n-1)}, \boldsymbol{\theta}^*\right)$ **then**

Accept the proposed parameter values and add to the chain: $\boldsymbol{\theta}^{(n)} = \boldsymbol{\theta}^*, \hat{J}^{(n)} = \hat{J}^*$.

**else**

Reject the proposed parameter values and keep the previous parameter values:

$\boldsymbol{\theta}^{(n)} = \boldsymbol{\theta}^{(n-1)}, \hat{J}^{(n)} = \hat{J}^{(n-1)}$.

Construct a sample of parameters from the posterior distribution and the corresponding log-likelihoods: $\left\{\left(\boldsymbol{\theta}^{(n)}\middle|\boldsymbol{y}\right); \hat{J}^{(n)}\right\}_{n=1}^{N}$.

### 3. Model specifications and corresponding likelihoods.

For analyzing the T cell lineage trees, we considered several model hypotheses about the diversification of single-cell-derived T cell populations into distinct subsets. We characterized different subsets with their distinct division speeds and modeled them as latent variables in our approach. We used a branching process framework (Harris, 1963) to model this diversification: we assumed that upon every cell division, each of the two daughter cells can (independently) either adopt the subset of the mother cell, or change into a different subset according to the model topology. Each of these choices can occur with probabilities that are part of the model parameters. In the following, we first define these model hypotheses, and then describe the simulation of latent variables and likelihood calculation for them.

**3.1. Model hypotheses tested in this study.** A schematic of the following models is shown in Figure 2C and 2J of the main text:

1. *Model #1.* We assume that no diversification occurs during the expansion of the T cell families and therefore the whole cell population follows the same inter-division time statistics. This "homogeneous" subset is denoted by $H$. The only model parameters are the mean and the CV of the inter-division time distribution: $\boldsymbol{\theta} = \{\mu_H, \mathrm{CV}_H\}$. Since all cells belong to one subset, there are no latent variables in this model.

2. *Model #2.* This model assumes that after an initial phase of semi-concordant divisions, the cells differentiate into another subset with different inter-division time statistics. The subset in the early expansion phase is referred to as Early-Activated ($EA$) and the subsequent subset is denoted by $H$. Upon every division of $EA$ cells, each of the daughter cells can either stay in the $EA$ subset with a certain probability or differentiate into $H$ otherwise. The model allows differentiation probabilities to vary between the phase 1 (subtree $T_0$ in Fig. S22) and phase 2 (subtrees $T_1, T_2, \dots, T_8$ in Fig. S22) of expansion; see Model #3 for the justification of this variation. The model parameters are the mean and CV of the inter-division time distribution of the subsets $EA$ and $H$ ($\mu_{EA}, \mathrm{CV}_{EA}, \mu_H, \mathrm{CV}_H$), and the probability of remaining in subset $EA$ in the two expansion phases ($p_{\mathrm{EA,1}}, p_{\mathrm{EA,2}}$): $\boldsymbol{\theta} = \{p_{\mathrm{EA,1}}, p_{\mathrm{EA,2}}, \mu_{EA}, \mathrm{CV}_{EA}, \mu_H, \mathrm{CV}_H\}$. The latent variable (i.e. subset) for a cell $m$ in model #2 can have two values: $x_m \in \{EA, H\}$.

3. *Model #3.* Similar to model #2, this model assumes an initial phase of semi-concordant divisions (subset $EA$). However, model #3 assumes a bifurcating pathway of diversification where the cells can differentiate out of $EA$ subset into two distinct subsets of fast and slowly dividing cells (subsets $F$ and $S$ respectively). This differentiation can happen at every $EA$ cell division with specific probabilities for staying in the $EA$ subset or differentiating into $F$ or $S$. In our T cell trees, the first few divisions are considerably less variable than the subsequent divisions (see the main text). To mimic this aspect, model #3 allows the differentiation probabilities to vary between the two expansion phases: the probability of differentiating from $EA$ into $S$ or $F$ in phase 2 (subtrees $T_1, T_2, \dots, T_8$) is constrained to be higher than 90%, while it is unconstrained in phase 1 (subtree $T_0$). The ratio between the differentiation probabilities into $S$ and $F$ subsets ($\mathrm{ratio}_{\mathrm{Slow}} = \frac{\text{differentiation probabilities into } S}{\text{differentiation probabilities into } F}$) is set to be the same in the two phases. To directly capture the distinction between the $F$ and $S$ subsets, the difference between their mean inter-division times is modeled as a parameter. The model parameters are the mean and CV of the inter-division time distribution of the $EA$ and $F$ subsets ($\mu_{EA}, \mathrm{CV}_{EA}, \mu_F, \mathrm{CV}_F$), the difference in the mean inter-

division time of subset $S$ w.r.t. subset $F$, and the CV of its inter-division time distribution ($d_{\mu_S} = \mu_S - \mu_F, \mathrm{CV}_S$), the probability of cells remaining in the $EA$ subset in the two expansion phases ($p_{\mathrm{EA},1}, p_{\mathrm{EA},2}$) and the ratio between the differentiation probabilities into $S$ and $F$ subsets ($\mathrm{ratio}_{\mathrm{Slow}}$): $\boldsymbol{\theta} = \left\{p_{\mathrm{EA},1}, p_{\mathrm{EA},2}, \mathrm{ratio}_{\mathrm{Slow}}, \mu_{EA}, \mathrm{CV}_{EA}, \mu_F, \mathrm{CV}_F, d_{\mu_S}, \mathrm{CV}_S\right\}$. The latent variable (i.e. subset) for a cell $m$ in this model can have three values: $x_m \in \{EA, S, F\}$.

4. Mixture model. This model assumes three subsets as in model #3. Initially cells belong to the $EA$ subset; upon every division, each daughter cell can stay in the $EA$ subset with a certain probability or change into a Mixture type ($M$) otherwise. The latter represents a mixture of two distinct subsets of fast and slowly dividing cells (subsets $F$ and $S$ respectively). Like model #3, different differentiation probabilities for the two expansion phases are assumed. The subset of $M$ cells is not certainly known; instead, it is assumed that they belong to subset $S$ with probability $\mathrm{ratio}_{\mathrm{Slow}}$ and to subset $F$ with probability $1 - \mathrm{ratio}_{\mathrm{Slow}}$. The model parameters are the mean and CV of the inter-division time distribution of the $EA$ and $F$ subsets ($\mu_{EA}, \mathrm{CV}_{EA}, \mu_F, \mathrm{CV}_F$), the difference in the mean inter-division time of subset $S$ w.r.t. subset $F$, and the CV of its inter-division time distribution ($d_{\mu_S}, \mathrm{CV}_S$), the probability of cells remaining in the $EA$ subset in the two expansion phases ($p_{\mathrm{EA},1}, p_{\mathrm{EA},2}$) and the probability that the cells in the mixture type possess subset $S$ ($\mathrm{ratio}_{\mathrm{Slow}}$): $\boldsymbol{\theta} = \left\{p_{\mathrm{EA},1}, p_{\mathrm{EA},2}, \mathrm{ratio}_{\mathrm{Slow}}, \mu_{EA}, \mathrm{CV}_{EA}, \mu_F, \mathrm{CV}_F, d_{\mu_S}, \mathrm{CV}_S\right\}$. The latent variable (i.e. type) for a cell $m$ can have two values: $x_m \in \{EA, M\}$.

   We emphasize that this "mixture model" does not intend to model an actual topology for the diversification of T cells. Rather, it provides a mathematical model alternative in which the same number of subsets and parameters as in model #3 is assumed. We compare this model against model #3 to test whether the superior performance of model #3 over models #1 and #2 is merely due to its additional flexibility, or whether the topological pathway encoded in model #3 is indeed essential.

5. *Model #4.* This model assumes the same diversification pathway as in model #3 and adds additional variability between different family trees (interfamily variation). It assumes that the mean inter-division time of subset $F$ is distributed across different trees according to a log-normal distribution with unknown mean and coefficient of variation ($\tilde{\mu}_F, \widetilde{\mathrm{CV}}_F$):

$$\mu_F \sim \log N\ \left(\tilde{\mu}_F, \widetilde{\mathrm{CV}}_F\right) \tag{3.1}$$

   It further assumes that the difference between the mean inter-division time of different subsets is constant in all trees. This implies that if subset $F$ in a tree is slower than the average, then subsets $EA$ and $S$ are also slower with the same distance to the average. In this way, all subset-specific mean inter-division times—for subsets $EA$, $F$ and $S$—are log-normally distributed across different trees. The model parameters are the mean and CV for the log-normal distribution of $\mu_F$ ($\tilde{\mu}_F, \widetilde{\mathrm{CV}}_F$), the differences in the mean inter-division time between the subsets ($d_{\mu_{EA}} = \mu_{EA} - \mu_F, d_{\mu_S} = \mu_S - \mu_F$), the subset-specific CVs of the inter-division time distribution ($\mathrm{CV}_{EA}, \mathrm{CV}_F, \mathrm{CV}_S$), and the differentiation probabilities as in model #3:

   $\boldsymbol{\theta} = \left\{p_{\mathrm{EA},1}, p_{\mathrm{EA},2}, \mathrm{ratio}_{\mathrm{Slow}}, \tilde{\mu}_F, \widetilde{\mathrm{CV}}_F, \mathrm{CV}_F, d_{\mu_{EA}}, \mathrm{CV}_{EA}, d_{\mu_S}, \mathrm{CV}_S\right\}$. The latent variable (i.e. subset) for a cell $m$ in model #4 can have three values: $x_m \in \{EA, S, F\}$.

**3.2. Simulation of the latent variables.** To calculate the likelihood of the data we need to simulate samples from the latent variables (Section 1.3.3). In model #1 all cells belong to the same subset and therefore no latent variables exist. For the rest of the models, we use the model topology and parameters to simulate the subsets of cells along a family tree. We assign the first founder cell of the tree to the Early-Activated subset as this subset is the starting point of all the considered pathways. We move along the family tree and upon every cell division randomly assign the subset of the daughter cells. Given the simulated subset of the mother cell, the possible subsets for each daughter cell are determined based on the model topology. The probability of a daughter cell to adopt any of these possible subsets is given by the model parameters (i.e. the differentiation probabilities). In this way, the tree structure and mother-daughter relationships are taken into account in simulating the latent variables.

For example in model #3, if a mother cell belongs to subset $EA$ in the first expansion phase, each daughter cell is independently assigned to subset $EA$ with probability $p_{EA,1}$, subset $S$ with probability $\mathrm{ratio}_{\mathrm{Slow}}(1 - p_{EA,1})$ and subset $F$ with probability $(1 - \mathrm{ratio}_{\mathrm{Slow}})(1 - p_{EA,1})$. However, if the mother cell belongs to either of the $S$ and $F$ subsets, both daughter cells directly inherit the subset of the mother cell.

**3.3. Calculation of the likelihood for models #1-4.** We need to calculate the likelihood of every cell given the latent variables and model parameters to obtain the overall data likelihood (see Equations 1.27 and 1.28 in Section 1.3.3). We assume that the inter-division of cells are log-normally distributed with subset-specific mean and coefficient of variation. If we denote the latent variable corresponding to cell $m$ by $x_m$, describing the subset to which cell $m$ belongs, the log-transformed inter-division time of cell $m$ is distributed according to the normal distribution $N\ (\mu_{x_m}, \mathrm{CV}_{x_m})$; $\mu_{x_m}$ and $\mathrm{CV}_{x_m}$ are determined by the model parameters. The likelihood of the lifetime of cell $m$, $t_m$, is then

$$\begin{cases} \pi(t_m|\mu_{x_m}, \mathrm{CV}_{x_m}) = \dfrac{1}{\mu_{x_m}\ \mathrm{CV}_{x_m}\sqrt{2\pi}}\ e^{-\frac{1}{2}\left(\frac{t_m - \mu_{x_m}}{\mu_{x_m}\ \mathrm{CV}_{x_m}}\right)^2}, & \text{if cell } m \text{ divided in the course of the experiment} \\ \pi(t_m|\mu_{x_m}, \mathrm{CV}_{x_m}) = 1 - \displaystyle\int_0^{t_m} \dfrac{1}{\mu_{x_m}\ \mathrm{CV}_{x_m}\sqrt{2\pi}}\ e^{-\frac{1}{2}\left(\frac{s - \mu_{x_m}}{\mu_{x_m}\ \mathrm{CV}_{x_m}}\right)^2} ds, & \text{if cell } m \text{ did not divide in the course of the experiment} \end{cases}$$

(3.2)

According to Eq. 3.2, if a cell has divided, its lifetime represents its inter-division time and therefore the PDF of the normal distribution $N\ (\mu_{x_m}, \mathrm{CV}_{x_m})$ is used to calculate its likelihood. But if a cell has not divided in the course of the experiment, its lifetime only provides a lower-bound for its inter-division time and therefore, the CDF of the corresponding normal distribution is used.

**3.4. Calculation of the likelihood for the mixture model.** In the mixture model, the subset of cells in the mixture type ($M$) is not directly known; instead it is assumed that they could belong to subset $S$ with probability $\mathrm{ratio}_{\mathrm{Slow}}$ and to subset $F$ with probability $1 - \mathrm{ratio}_{\mathrm{Slow}}$. Therefore Eq. 3.2, that assumes the subset of cells is known given the latent variables, cannot be used for likelihood calculation. Instead, we use the following equation for cells of type $M$ based on the likelihood for a mixture model (Pyne et al., 2009):

$$
\begin{cases}
\pi(t_m) = \text{ratio}_{\text{Slow}} \dfrac{1}{\mu_S\ \text{CV}_S \sqrt{2\pi}}\ e^{-\frac{1}{2}\left(\frac{t_m - \mu_S}{\mu_S\ \text{CV}_S}\right)^2} + \\
(1 - \text{ratio}_{\text{Slow}})\ \dfrac{1}{\mu_F\ \text{CV}_F \sqrt{2\pi}}\ e^{-\frac{1}{2}\left(\frac{t_m - \mu_F}{\mu_F\ \text{CV}_F}\right)^2}, & \text{if cell } m \text{ divided in the course of the experiment} \\
\pi(t_m) = \text{ratio}_{\text{Slow}} \left(1 - \displaystyle\int_0^{t_m} \dfrac{1}{\mu_S\ \text{CV}_S \sqrt{2\pi}}\ e^{-\frac{1}{2}\left(\frac{s - \mu_S}{\mu_S\ \text{CV}_S}\right)^2} ds\right) + \\
(1 - \text{ratio}_{\text{Slow}}) \left(1 - \displaystyle\int_0^{t_m} \dfrac{1}{\mu_F\ \text{CV}_F \sqrt{2\pi}}\ e^{-\frac{1}{2}\left(\frac{s - \mu_F}{\mu_F\ \text{CV}_F}\right)^2} ds\right), & \text{if cell } m \text{ did not divide in the course of the experiment}
\end{cases}
$$

(3.3)

Equation 3.3 describes the likelihood of the lifetime of a cell as the weighted sum of the likelihoods if it belonged to subsets $S$ and $F$ respectively. For cells in the $EA$ subset, Eq. 3.2 is used to calculate their likelihood.

**3.5. Inference results for different models.** We split our data into eight groups of five trees and performed parameter inference and model comparison on all eight data groups. The inferred parameter posteriors, model evidences and Bayes factors for every data group are shown in Figures S4, S6, S7, S12, S13 and S17-19. For better comparability, we reported base-10 logarithm of model evidences and Bayes factors. In addition to the group-wise metrics, we calculated average model evidences and Bayes factors from the results of all data groups together. The average log10-model evidence was calculated as $\frac{1}{8}\sum_{i=1}^{8} ME_i$, where $ME_i$ is the median of the log10-model evidence calculated in data group $i$. The average log10-Bayes factor was calculated as $\frac{1}{8}\sum_{i=1}^{8} BF_i$, where $BF_i$ is the median of the log10-Bayes factor calculated in data group $i$. The average model evidences and Bayes factors are shown in Figures 2D, 2L, S3 and S5.

### 4. Statistical features of the experimental data.

**4.1. Contribution of intrafamily and interfamily variability in the overall variation of inter-division times.** We calculated the total variance of the inter-division times and the contribution of intrafamily and interfamily sources in the experimental data. Intrafamily variance was calculated as the weighted mean of the variances of inter-division times within different families:

$$
\text{Intrafamily variance} = \frac{\sum_{i=1}^{n_{\text{tree}}} n_{\text{cell}_i}\, \sigma_i^2}{\sum_{i=1}^{n_{\text{tree}}} n_{\text{cell}_i}} \tag{4.1}
$$

where $n_{\text{cell}_i}$ and $\sigma_i^2$ are respectively the number of cells and the variance of inter-division times in the $i^{\text{th}}$ tree. Interfamily variance was calculated as the weighted variance of the family mean inter-division times:

$$
\text{Interfamily variance} = \frac{\sum_{i=1}^{n_{\text{tree}}} n_{\text{cell}_i}\, (\mu_i - \mu^*)^2}{\sum_{i=1}^{n_{\text{tree}}} n_{\text{cell}_i} - 1}\ , \quad \mu^* = \frac{\sum_{i=1}^{n_{\text{tree}}} n_{\text{cell}_i}\, \mu_i}{\sum_{i=1}^{n_{\text{tree}}} n_{\text{cell}_i}} \tag{4.2}
$$

where $\mu_i$ is the mean inter-division time in the $i^{\text{th}}$ tree, and $\mu^*$ is the weighted mean of $\mu_i$ for $i = 1, \dots, n_{\text{tree}}$. These variance terms for the experimental data are shown in Figure 1G.

**4.2. Percentage of the trees whose four branches have significantly distinct inter-division times.** For every tree in the experimental data, we compared the inter-division time distribution of the four branches of the tree as depicted in Figure 1I. We performed one-way ANOVA to test whether the four branches have different mean inter-division times. We then estimated positive false discovery rates (pFDR) for multiple hypothesis testing based on the Benjamini and Hochberg method (Benjamini and Hochberg, 1995) across all trees. We selected the trees that had a pFDR smaller than 0.05 as trees whose four branches have significantly distinct inter-division times. These selected trees are highlighted in Figure 1J.

### 5. Simulation studies for model validation.

To inspect how well the studied models were matching to the experimental data—beyond measures provided by model evidence and Bayes factors—we performed the following simulation studies. We examined how well the simulated data based on different models captured the statistical features of the experimental data.

**5.1. Overall distribution of the inter-division times.** We simulated 10,000 family trees based on the model topologies #3 and #4, and the inferred posterior distribution for the corresponding model parameters. We obtained the distribution of the inter-division times of all cells regardless of their subset from all simulated trees together; we compared this against the distribution of inter-division times in the experimental data. We further obtained the distribution of inter-division times of cells in the subsets $EA$, $S$ and $F$ in the simulated data for illustration purposes. The results are shown in Figures S8 and S14 for model #3 and model #4 respectively.

**5.2. Contribution of intrafamily and interfamily variability in the overall variation of inter-division times.** We simulated 500 datasets of each 44 trees based on model topologies #3 and #4, and the inferred posterior distribution for the corresponding model parameters. For each dataset, we calculated the total variance of the inter-division times and the contribution of intrafamily and interfamily sources. Intrafamily variance was calculated as the weighted mean of the variances of inter-division times within different families:

$$\text{Intrafamily variance} = \frac{\sum_{i=1}^{44} \hat{n}_i \, \hat{\sigma}_i^2}{\sum_{i=1}^{44} \hat{n}_i} \tag{5.1}$$

where $\hat{n}_i$ and $\hat{\sigma}_i^2$ are respectively the number of cells and the variance of inter-division times in the $i^{\text{th}}$ simulated tree. Interfamily variance was calculated as the weighted variance of the family mean inter-division times:

$$\text{Interfamily variance} = \frac{\sum_{i=1}^{44} \hat{n}_i \, (\hat{\mu}_i - \bar{\mu})^2}{\sum_{i=1}^{44} \hat{n}_i - 1} \;, \quad \bar{\mu} = \frac{\sum_{i=1}^{44} \hat{n}_i \, \hat{\mu}_i}{\sum_{i=1}^{44} \hat{n}_i} \tag{5.2}$$

where $\hat{\mu}_i$ is the mean inter-division time in the $i^{\text{th}}$ simulated tree, and $\bar{\mu}$ is the weighted mean of $\hat{\mu}_i$ for $i = 1, \dots, 44$. We then obtained the 90%-confidence interval for the total variance, intrafamily variance and interfamily variance in the 500 simulated datasets. We examined whether these confidence intervals contained the corresponding variance terms calculated for the experimental data. The results for the simulations based on model #3 and model #4 are shown in Figures S9 and S15 respectively.

**5.3. Percentage of the trees whose four branches have significantly distinct inter-division times.** We used the same simulated data as in Section 5.2. For every tree in a simulated dataset, we compared the inter-division time distribution of the four branches of the tree as depicted in Figure 1I. We performed one-way

ANOVA to test whether the four branches have different mean inter-division times. We then estimated positive false discovery rates (pFDR) for multiple hypothesis testing based on the Benjamini and Hochberg method (Benjamini and Hochberg, 1995) across all 44 trees in the dataset. We selected the trees that had a pFDR smaller than 0.05 as trees whose four branches have significantly distinct inter-division times, and in every dataset calculated the percentage of these selected trees. We then obtained the distribution of this percentage across the 500 simulated datasets and examined whether this distribution contained the percentage calculated from the experimental data. The results for the simulations based on models #3 and #4 are shown in Figures S10 and S16 respectively.

### References:


1. Benjamini Y, Hochberg Y (1995) Controlling the false discovery rate: A practical and powerful approach to multiple testing. *J. Royal Stat. Soc*. 57:289–300.

2. Harris T (1963) *The Theory of Branching Processes*. Springer, Berlin.

3. Hilsenbeck O, et al. (2016) Software tools for single-cell tracking and quantification of cellular and molecular properties. *Nat Biotechnol* 34(7):703–706.

4. Loos C (2016) *Analysis of single-cell data: ODE-constrained mixture modeling and approximate Bayesian computation.* Best Masters. Springer.

5. Loos C, et al. (2016) Parameter Estimation for Reaction Rate Equation Constrained Mixture Models. In: Bartocci E., Lio P., Paoletti N. (eds) *Computational Methods in Systems Biology*. CMSB 2016. Lecture Notes in Computer Science, vol 9859. Springer, Cham.

6. Pyne S, et al. (2009) Automated high-dimensional flow cytometric data analysis. *Proc. Natl. Acad. Sci.* USA, 106, 8519–8124.

7. Vyshemirsky V, Girolami MA (2008) Bayesian ranking of biochemical system models. *Bioinformatics*, 24(6):833–839.

8. Wilkinson DJ (2009) Stochastic modelling for quantitative description of heterogeneous biological systems. *Nat. Rev. Genet.* 10:122-133.

9. Wilkinson DJ (2011) *Stochastic Modelling for Systems Biology*. Second Edition. CRC Press.
